# Beyond consensus sequence: a quantitative scheme for inferring transmission using deep sequencing in a bacterial transmission model

**DOI:** 10.1101/2022.10.17.512634

**Authors:** Madikay Senghore, Hannah Read, Priyali Oza, Sarah Johnson, Hemanoel Passarelli-Araujo, Bradford P Taylor, Stephen Ashley, Alex Grey, Alanna Callendrello, Robyn Lee, Matthew R Goddard, Thomas Lumley, William P Hanage, Siouxsie Wiles

## Abstract

Genomic surveillance provides a data source complementary to contact tracing to resolve putative transmission chains. However, the role of within-host diversity in transmission is understudied due to a lack of experimental and clinical datasets that capture within-host diversity in both donors and recipients. Here, we assess the utility of deep-sequenced genomic surveillance within a mouse transmission model where the gastrointestinal pathogen *Citrobacter rodentium* was controllably spread during co-housing of infected and naïve animals. We observed that within-host variants were maintained over multiple transmission steps until fixation or elimination. We present a model for inferring the likelihood that a given pair of samples are linked by transmission, by comparing the allelic frequency at variant genomic *loci*. Our data affirm that within-host single nucleotide variants (iSNVs) can repeatedly pass from donor to recipient along the transmission chain, and the mere sharing of iSNVs between different transmission pairs offers limited confidence in identifying a transmission pair. Beyond the presence and absence of within-host variants, we show that differences arising in the relative abundance of iSNVs can infer transmission pairs with high precision. An important component of our approach is that the inference is based solely on sequence data, without incorporating epidemiological or demographic data for context. Our model, which substantially reduces the number of comparisons a contact tracer needs to consider, may enhance the accuracy of contact tracing and other epidemiological processes, including early detection of emerging transmission clusters.

## Introduction

The control and/or elimination of infectious diseases depend on identifying and interrupting transmission chains, particularly during acute epidemics ^1–3^. While classical epidemiological techniques such as contact tracing remain integral to the epidemic control toolkit, they can be supported and strengthened by modern technological approaches ^4^. For example, genomic analysis can shed light on the emergence of variants to reconstruct transmission chains in an epidemic ^5–9^. The increased resolution provided by genomics has allowed epidemiologists to guide both reactive measures through retracing the transmission routes in outbreaks ^6^ and proactive measures after discovering environmental reservoirs ^10^.

Whole genome sequencing is also an essential tool to identify novel routes of host-to-host transmission^11^. However, using genomics to infer transmission is limited by the diversity of the circulating pathogen population; two genetically similar bacteria may indicate genuine transmission between hosts, may be a result of different introductions from a third host, or arise from independent lineages with the same de novo mutational events (but with increasing numbers of unique differences this become extremely improbable). This limit is especially acute during a rapidly spreading epidemic where transmission occurs faster than fixed mutations are accumulated.

The next frontier in genomic surveillance is moving beyond single nucleotide variants (SNVs) based on consensus sequences to include further variation uncovered through deep sequencing. This allows the identification of within-host single nucleotide variants (iSNVs) present at frequencies lower than the threshold set to define a fixed mutation. The sharing of iSNVs in two closely related isolates provides extra information to aid the inference of transmission chains ^7,12,13^. As *de novo* mutations arise, they can be maintained and passed on during transmission ^6,8,14^. When transmission occurs, a bottleneck modulates how SNVs and iSNVs are passed on from the donor to the recipient ^14–16^. Estimating bottleneck sizes is challenging, but by considering stochasticity in the donor and recipient, it is possible to infer bottleneck sizes from deep sequence data ^15,16^.

We have previously created a bioluminescent derivative of the mouse enteropathogen *Citrobacter rodentium* ^17,18^ to track the bacterium within infected mice and the environment ^19– 22^. Here, we established ten transmission chains where *C. rodentium* ICC180 was controllably spread during mouse co-housing; that is, we know with certainty who infected whom at each stage. We deep sequenced the bacteria shed from each infected animal at the point of transmission. Moreover, we introduce further methods to aid identification of transmission pairs by quantifying differences in the allelic frequency at iSNVs and SNV *loci* between sample pairs. Here we test the hypothesis that within host diversity can be used to offer a quantitative measure showing which isolates pairs are more likely to be linked by transmission events. Our approach provides an accurate, quantifiable, and actionable measure that can enhance contact tracing and inference of transmission chains, if done in a timely manner.

## Results

### Observed dynamics of the transmission model in mice

Throughout this study, 220 mice were infected with/exposed to *C. rodentium* ICC180 within 10 transmission chains (Figure 1, Supplementary Table 1). Using linear mixed models, we observed no statistically significant differences in weight losses or gains between chains or between treatments (with or without nalidixic acid supplementation) (P value ⋝ 0.05). We monitored transmission and infection dynamics by measuring viable bacterial counts (Figure 2A-C) and bioluminescence from shed stool and by non-invasive biophotonic imaging (Figure 1D). We observed no changes in the anatomical location of the infecting bacteria (Figure 1D) or in disease severity, suggesting that the pathogenicity and disease dynamics of *C. rodentium* ICC180 remained unchanged over the course of the experiment. Of the animals who successfully infected their cage-mates, infections progressed as expected, with a rapid increase in bacterial numbers within the first few days, followed by a peak/plateau, and then a decline (Figure 1A).

**Figure 1.**
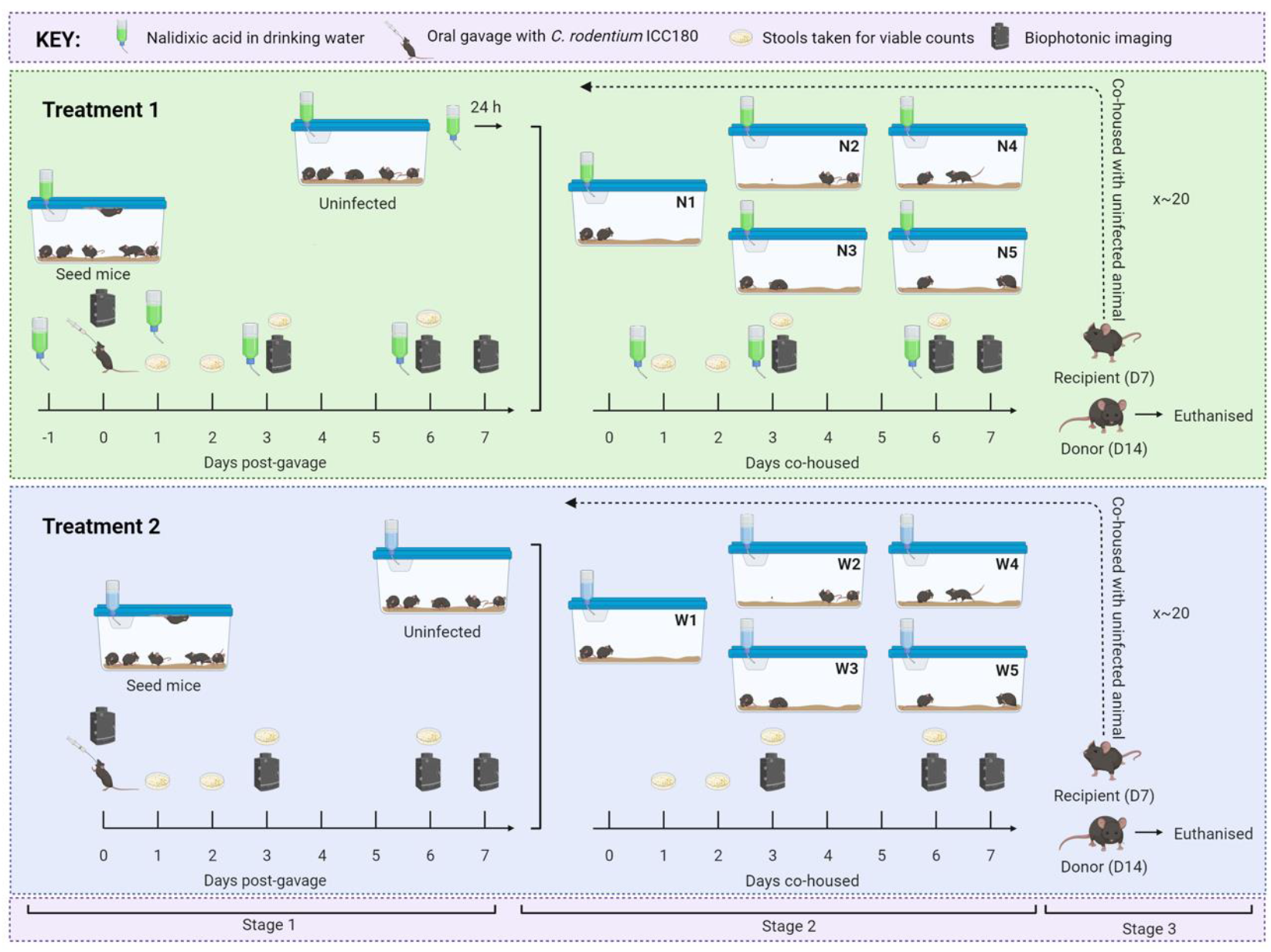
Experimental schematic summarizing the establishment of ten mouse-to-mouse *Citrobacter rodentium* ICC180 transmission chains. Seed mice were split into two treatment groups (with nalidixic acid added to the drinking water every 2-3 days [N] or without [W]) and orally gavaged with ICC180. Seven days post-gavage, donor animals were transferred to individual cages (N1-5/W1-5) and cohoused with an uninfected cage-mate (recipient). After seven days, the donor was humanely euthanized. The recipient then became the donor for the next step in the transmission chain by being transferred to a clean cage and cohoused with an uninfected cage-mate. This cycle was repeated until the end of the experiment. Infection and transmission dynamics were monitored by measuring luminescence and viable bacterial counts from stool samples and in vivo by biophotonic imaging.

**Figure 2.**
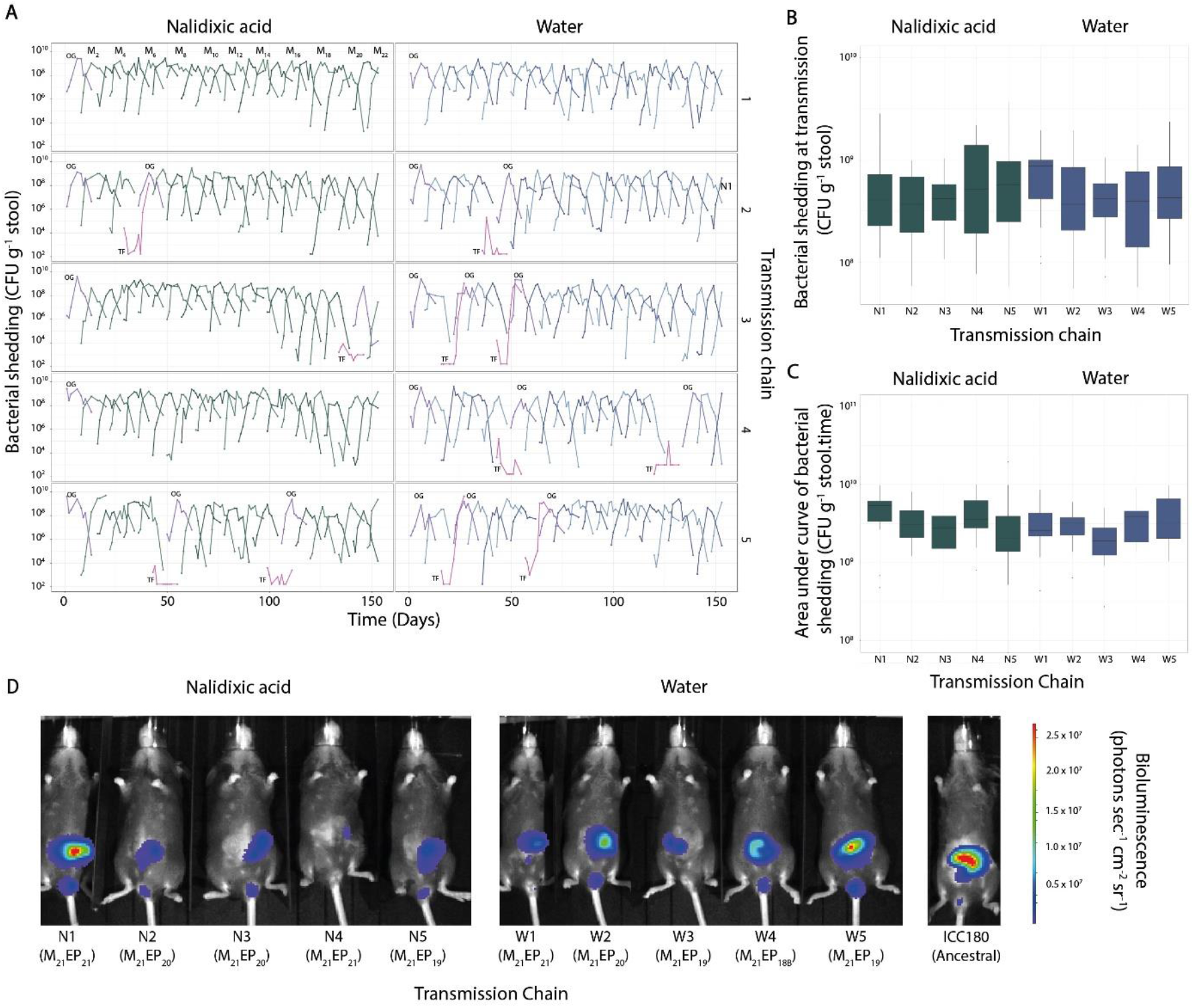
*In vivo* infection and transmission dynamics of 10 mouse-to-mouse *C. rodentium* ICC180 transmission chains. **A)** Bacterial shedding in stool of each animal (as colony forming units [CFU]^−1^ stool) over 14 days. Animals infected by oral gavage (OG) are shown in purple, while those who failed to be infected by natural transmission (TF, transmission failure) are shown in pink. **B)** Box-whisker plot of bacterial shedding (as CFU^−1^ stool) at transmission summarised by transmission chain. **C)** Box-whisker plot of the area under curve values of bacterial shedding (as CFU^−1^ stool.time) by transmission chain. **D)** *In vivo* location of *C. rodentium* ICC180 within M_21_ donor mice at the time of transmission compared to ancestral ICC180. Key: OG, oral gavage; TF, transmission failure.

We also observed a large variation in the number of viable *C. rodentium* ICC180 shed by each animal around the time of transmission (Figure 2B, Table 1), ranging from 6.83 × 10^6^ colony forming units (CFU) g^−1^ stool (W3) to 5.33 × 10^9^ CFU g^−1^ stool (W2) (Table 1). Bacterial numbers were lower in animals without nalidixic acid supplementation, but with low statistical support (Table 2). However, the transmission chains differed in both viable counts and *in vivo* bioluminescence data, suggesting that each chain behaved independently (Table 3). Three transmission chains (N1, N4, and W1) experienced no transmission failures, while 11 animals from the remaining chains failed to become infected (Figure 2A, Table 1). These failures most likely reflect differences in animal grooming and coprophagic behavior. However, it is worth noting that three of the animals came from the same cohort of 10 (M_7_) and may have been littermates.

**Table 1.**
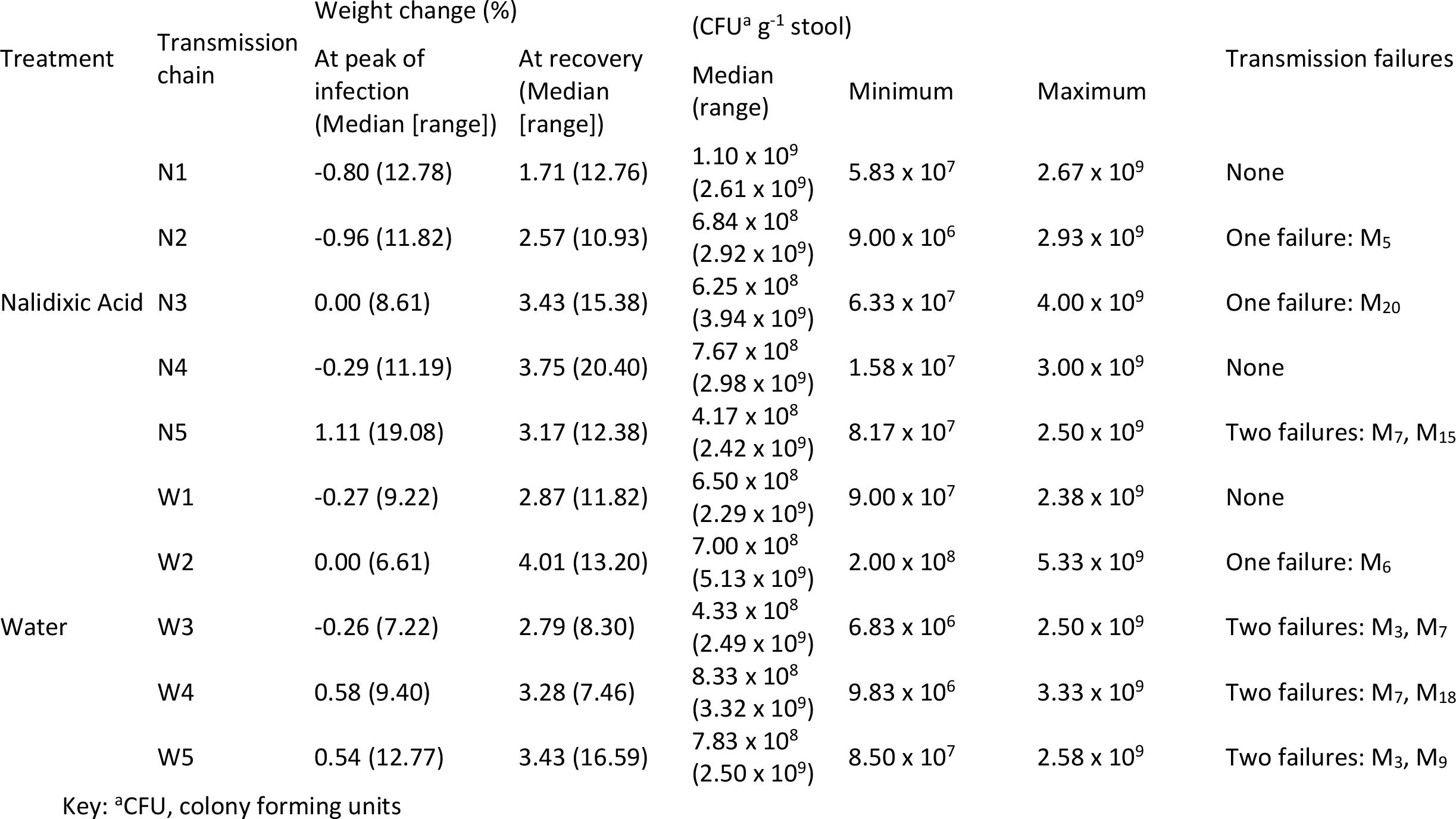
Summary of changes in animal weight, bacterial shedding prior to housing, and transmission failures, by transmission chain Viable counts prior to co-housing

| Treatment | Transmission chain | Weight change (%) |  | Viable counts prior to co-housing (CFU<sup>a</sup> g<sup>-1</sup> stool) |  |  | Transmission failures |
| --- | --- | --- | --- | --- | --- | --- | --- |
|  |  | At peak of infection (Median [range]) | At recovery (Median [range]) | Median (range) | Minimum | Maximum |  |
| Nalidixic Acid | N1 | -0.80 (12.78) | 1.71 (12.76) | 1.10 x 10<sup>9</sup> (2.61 x 10<sup>9</sup>) | 5.83 x 10<sup>7</sup> | 2.67 x 10<sup>9</sup> | None |
|  | N2 | -0.96 (11.82) | 2.57 (10.93) | 6.84 x 10<sup>8</sup> (2.92 x 10<sup>9</sup>) | 9.00 x 10<sup>6</sup> | 2.93 x 10<sup>9</sup> | One failure: M<sub>5</sub> |
|  | N3 | 0.00 (8.61) | 3.43 (15.38) | 6.25 x 10<sup>8</sup> (3.94 x 10<sup>9</sup>) | 6.33 x 10<sup>7</sup> | 4.00 x 10<sup>9</sup> | One failure: M<sub>20</sub> |
|  | N4 | -0.29 (11.19) | 3.75 (20.40) | 7.67 x 10<sup>8</sup> (2.98 x 10<sup>9</sup>) | 1.58 x 10<sup>7</sup> | 3.00 x 10<sup>9</sup> | None |
|  | N5 | 1.11 (19.08) | 3.17 (12.38) | 4.17 x 10<sup>8</sup> (2.42 x 10<sup>9</sup>) | 8.17 x 10<sup>7</sup> | 2.50 x 10<sup>9</sup> | Two failures: M<sub>7</sub>, M<sub>15</sub> |
| Water | W1 | -0.27 (9.22) | 2.87 (11.82) | 6.50 x 10<sup>8</sup> (2.29 x 10<sup>9</sup>) | 9.00 x 10<sup>7</sup> | 2.38 x 10<sup>9</sup> | None |
|  | W2 | 0.00 (6.61) | 4.01 (13.20) | 7.00 x 10<sup>8</sup> (5.13 x 10<sup>9</sup>) | 2.00 x 10<sup>8</sup> | 5.33 x 10<sup>9</sup> | One failure: M<sub>6</sub> |
|  | W3 | -0.26 (7.22) | 2.79 (8.30) | 4.33 x 10<sup>8</sup> (2.49 x 10<sup>9</sup>) | 6.83 x 10<sup>6</sup> | 2.50 x 10<sup>9</sup> | Two failures: M<sub>3</sub>, M<sub>7</sub> |
|  | W4 | 0.58 (9.40) | 3.28 (7.46) | 8.33 x 10<sup>8</sup> (3.32 x 10<sup>9</sup>) | 9.83 x 10<sup>6</sup> | 3.33 x 10<sup>9</sup> | Two failures: M<sub>7</sub>, M<sub>18</sub> |
|  | W5 | 0.54 (12.77) | 3.43 (16.59) | 7.83 x 10<sup>8</sup> (2.50 x 10<sup>9</sup>) | 8.50 x 10<sup>7</sup> | 2.58 x 10<sup>9</sup> | Two failures: M<sub>3</sub>, M<sub>9</sub> |

Key: <sup>a</sup>CFU, colony forming units

**Table 2.** Average differences in log bacterial measures (shedding and in vivo bioluminescence) per passage in a chain and between conditions (with or without nalidixic acid supplementation).

|  |  | Bacterial shedding |  | In vivo bioluminescence<br>(photons s<sup>-1</sup>) |  |
| --- | --- | --- | --- | --- | --- |
|  |  | CFU g<sup>-1</sup> stool | RLU g<sup>-1</sup> stool | Abdomen | Rectum |
| Step in chain | Log ratio | -0.093 | -0.016 | -0.024 | -0.001 |
|  | Standard deviation | 0.014 | 0.010 | 0.008 | 0.007 |
| Treatment | Log ratio | -0.379 | -0.379 | -0.137 | -0.264 |
|  | Standard deviation | 0.317 | 0.256 | 0.179 | 0.173 |

**Table 3.** Estimated standard deviation of each measure, on the log scale, between transmission chains and between mice within a chain, and the residual standard deviations.

|  | Bacterial shedding |  | In vivo bioluminescence<br>(photons s<sup>-1</sup>) |  |
| --- | --- | --- | --- | --- |
|  | CFU g<sup>-1</sup> stool | RLU g<sup>-1</sup> stool | Abdomen | Rectum |
| Mouse | 0.22 | 0.00 | 0.00 | 0.00 |
| Chain | 0.44 | 0.37 | 0.24 | 0.24 |
| Residual | 3.27 | 2.40 | 1.37 | 1.28 |

### Within host variants were transferred over successive transmission steps until they became fixed mutations

Our first objective was to keep track of the rate at which fixed mutations were being accrued over successive transmission steps. Fixed mutations were observed at 12 individual *loci* across the genome; the consensus sequences based on these 12 *loci* were used to reconstruct a phylogenetic tree (Figure 3A). We observed that about 41 isolates at the base of the tree had identical consensus sequences and did not possess any within-host variants (Figure 3A). On average, approximately one new within-host variant (iSNV) emerged with every transmission step, regardless of whether it went on to fixation, which translated to roughly 44 new iSNVs per genome per year (Figure 3B). On average, it took 31 days for an emergent iSNV to reach fixation, either through genetic drift or selection (median 14 days, range: 7 to 126 days) (Figure S1), which translated to 0.09 SNVs becoming fixed for every transmission step or approximately 5 SNVs per genome per year (Figure 3C). Across the entire dataset, the mean pairwise genetic distance between isolates was 1.05 SNVs (Figure 3D). 203 out of 205 isolates had an identical consensus sequence with at least one other isolate, including 126 isolates with the index strain’s consensus at the basal branch on the phylogeny (Figure 3A). Typically, *loci* emerged as iSNVs and fixed over multiple transmission steps, while other iSNVs drifted and eventually faded away (Figure 3E). Allelic frequency of *loci* changed throughout transmission chains including sweeps by subpopulations, signs of competing subpopulations and potential linkage between two mutations (Figure 3E). Both stochastic and selective processes will be operating on the various bacterial lineages, but we cannot disentangle these.

**Figure 3.**
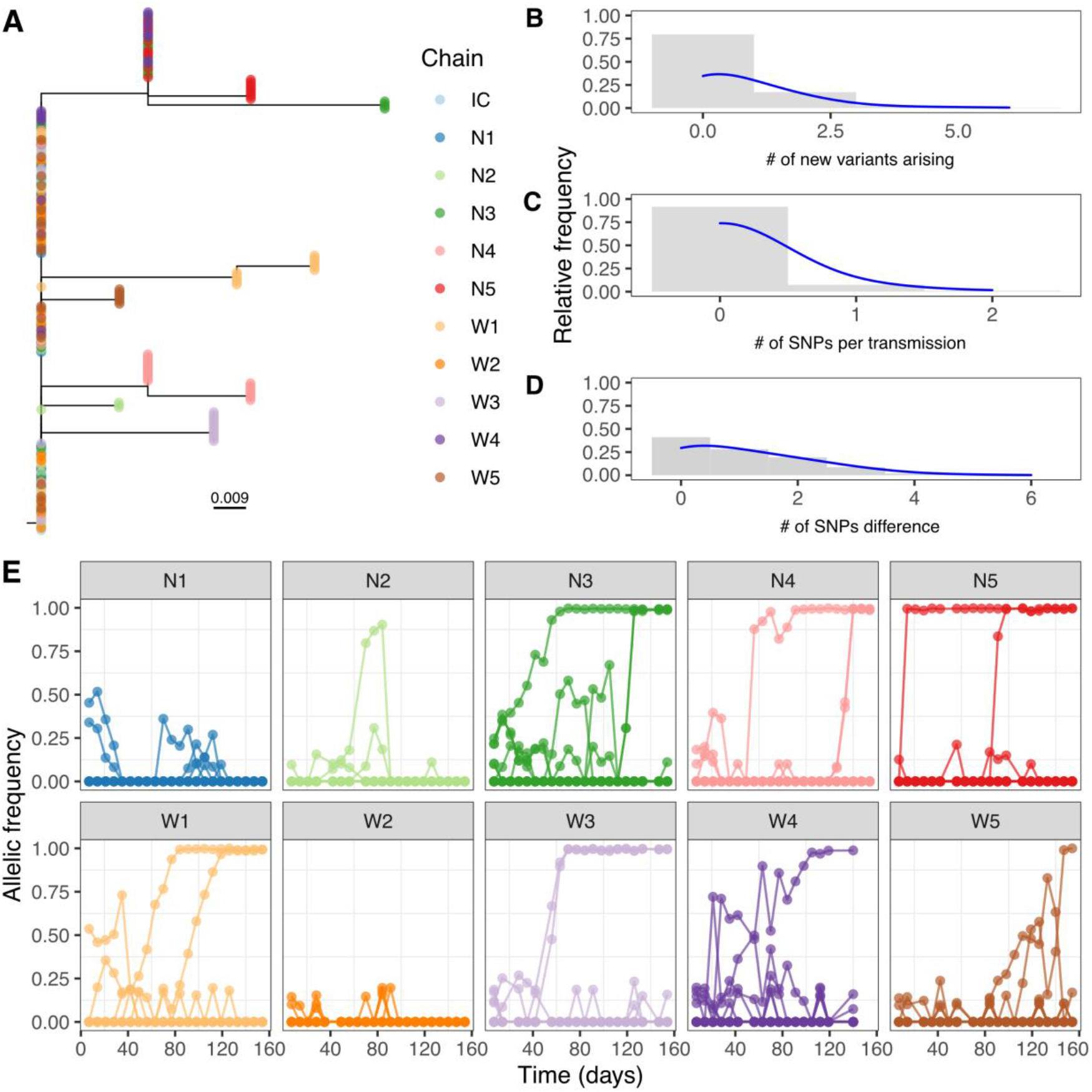
Accumulation of diversity along transmission chains. A) Maximum likelihood phylogenetic tree based on core SNPs with tips annotated by transmission chain. The “IC” chain refers to the reference ICC180, which was used as an outgroup to root the tree. B-D) Histograms showing the relative frequency for the number of iSNVs emerging in a transmission step, the number of SNVs becoming fixed during transmission, and the SNP distances across the full dataset, respectively. B) Line graph tracking the allelic frequency of each locus throughout the 10 transmission chains.

### Changes in the allele frequency can improve identification of transmission pairs

To distinguish strains beyond the consensus sequence, we recorded the allelic frequency of each iSNV and SNV and quantified the magnitude of the difference at these sites. The sum of all changes in the allelic frequency was divided by the number of sites where there was a difference, and this metric was referred to as the mean change in allelic frequency (per variable site). We noted that over successive transmission steps, the mean difference in allelic frequency increased with the number of transmission steps (Figure 4A). Moreover, linear regression showed that the mean difference in allelic frequency increases by approximately 0.02 units per transmission step (P-value < 0.001, adjusted R2 = 0.98) (Figure 4B).

**Figure 4.**
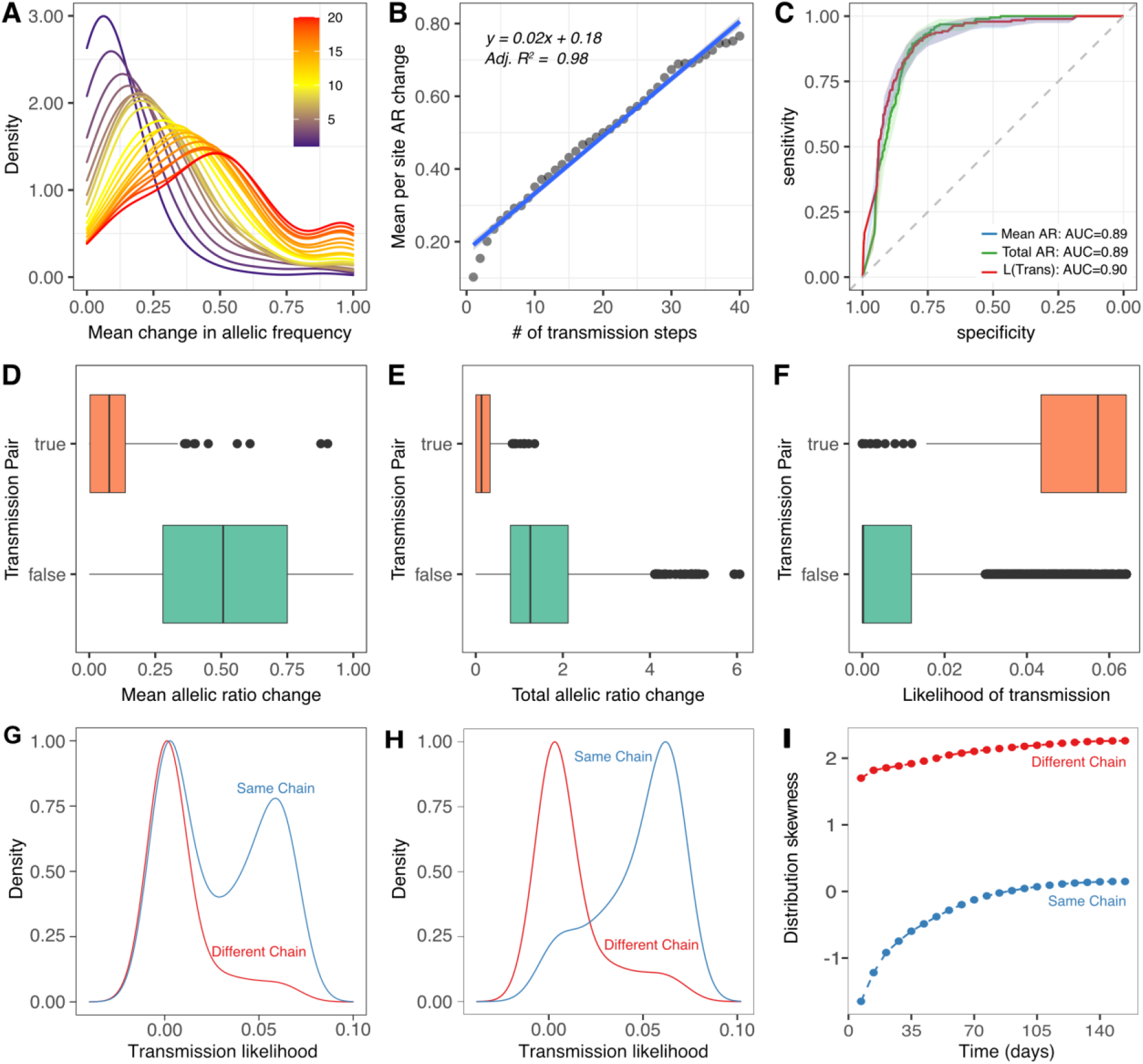
Allelic frequency at variable *loci* over successive transmission steps. A) Density distribution of mean allelic frequency across different transmission steps B) Linear plot of average of mean allelic frequency changes, grouped by number transmission C) ROC curves showing the performance of mean change in allelic frequency, total change in allele frequency, and the transmission likelihood as a predictor of transmission. D-F) Boxplots comparing the distribution of each metric in transmission pairs vs non transmission pairs. F) Distribution of transmission likelihoods between the same chain and in different chains considering donors in preceding transmission steps. G) Distribution of transmission likelihoods considering donors at maximum three precedent transmission steps. H) Skewness of the distribution based on incremental time / transmission steps (days).

We also explored how predictable transmission pairs were based on the allelic frequency change and the likelihood of transmission. The likelihood was obtained by using a Bayesian framework (see Methods for details). We plotted area under curves (AUC) for three different metrics (mean allelic frequency change, total allelic frequency change, and likelihood of transmission) to test how well they performed on distinguishing transmission pairs from non-transmission pairs when the cut off was varied. All these three metrics achieved an excellent performance on distinguishing transmission pairs (AUC > 0.89) (Figure 4C). Moreover, the mean and total changes in allelic ratio were lower in transmission pairs (P-value < 0.05, Mann Whitney U Test) (Figure 4D-E).

For each isolate, the inferred transmission likelihoods were used to identify the isolates most likely to be linked by transmission. Isolate pairs within the same transmission chain had a significantly higher transmission likelihood than isolates from different transmission chains (P-value < 0.05, Mann Whitney U Test) (Figure 4F). In addition to investigating the potential of transmission likelihoods to discriminate transmission pairs, we also evaluated how well this metric discriminates transmission chains, by considering potential donors in transmission steps before the recipient. For example, when considering mouse M_4_ in chain W5 we only considered mice from three prior transmission steps across all chains. By using only this assumption, we observed a bimodal distribution of likelihood of transmission for individuals within the same chain and a right-skewed distribution for those on different chains (Figure 4G). Therefore, high likelihood values indicate a greater propensity to detect isolates recovered from the same chain. Moreover, by restricting to three prior transmission steps, we observed that both distributions become oppositely asymmetric, increasing the ability to correctly predict the transmission chain only with the likelihood value (Figure 4H). Finally, the smaller the number of transmission steps (time since infection), the more left-skewed the distribution is (Figure 4I).

We also explored the potential of using inferred likelihood to reconstruct transmission chains (Figure 5), and to identify when the bacterium failed to transmit from mouse to mouse (Figure 5A). The reconstruction of the network based on likelihood had an excellent performance at detecting transmission clusters (Figure 5B). Interestingly, our model is also capable of identifying transmission at failure points such as in chain W3 where mouse M_4_ received the *C. rodentium* inoculum by oral gavage (see the break in the W3 strand in Figure 5, as highlighted in Figure 5B).

**Figure 5.**
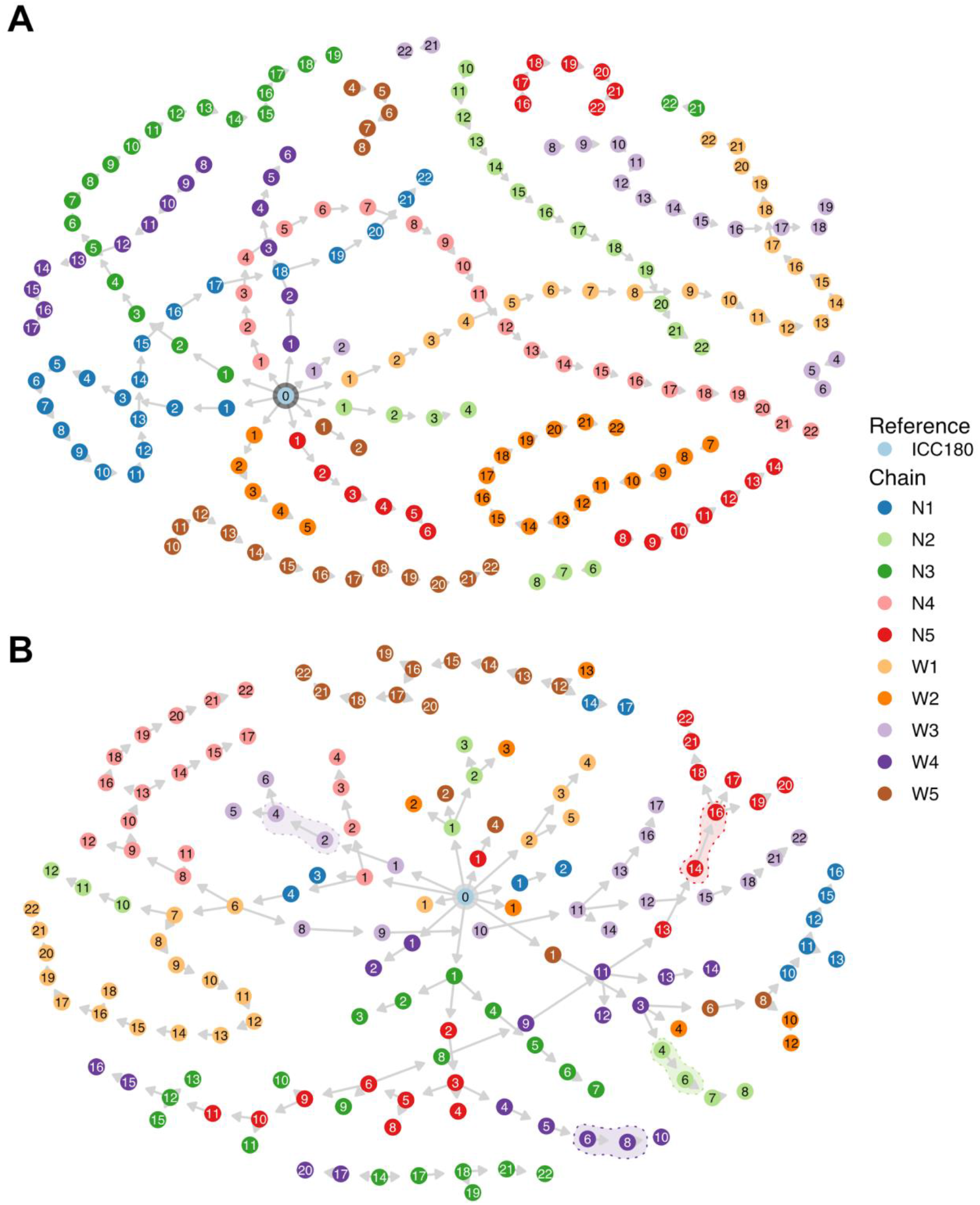
Reconstruction of the transmission network based on the inferred likelihood. A) ground of truth of transmissions from ICC180, including transmission failures (chain breaks). Colors indicate transmission chains. B) Transmission network from inferred likelihood values, using the time of infection as a *prior*. Transmission failures that were recognized by our method are highlighted.

To ensure that the observed mean changes in allelic frequency reflected actual differences arising over a transmission step, the bottleneck sizes for all transmission steps were estimated using the beta-binomial model. Confidence intervals (95% CI) for bottleneck size ranged from one to 32 and estimates above 32 were typically from pairs with a small mean change in allelic frequency. Bottleneck estimates above 32 showed progressively wider confidence intervals (Figure S2). Wider intervals were due to comparisons involving samples that had little or no within host diversity to serve as a basis for comparison.

### Simply sharing within host variants is insufficient for predicting transmission

We quantified how well the presence of within-host variants (allelic frequency > 0.025) in two isolates could predict whether two isolates were of a transmission pair or the same transmission cluster. On average, strains that were separated by more transmission steps shared fewer variants but there was notable overlap in the distributions (Supplementary Fig 4A). Although the mean number of shared iSNVs/ SNVs decreased monotonically with the number of transmission steps on average (Supplementary Figure 4B), this was insufficient to discriminate transmission pairs from other closely related pairs. Only some rarely observed variants (present in <5% of the study population) could predict transmission if they were shared by two isolates (Supplementary Fig 4C). Often, these rare variants persisted across multiple samples making them better predictors of whether a secondary isolate was within 5 or fewer transmission steps (Supplementary Figure 3D) and part of the same transmission chain (Supplementary Fig 4E).

## Discussion

Using shared variants to infer the transmission link between isolates, either directly or indirectly through intermediary transmission steps, is well supported ^7,12–14,23–25^. However, while shared variants indicate transmission, they do not provide a quantitative measure of who is most likely to have infected whom, especially when the variant is shared across multiple individuals. Previous works relied on simulated datasets to quantify the role of shared variants or employed Bayesian phylodynamic models to account for unsampled hosts ^7,26,27^. The challenge is that these methods are highly technical and do not always provide a quantifiable measure to inform non-bioinformatician public health officials. Our approach is unique because it quantifies the likelihood of two isolates being transmission pairs, prior to incorporating epidemiological and demographic data. Moreover, our method can differentiate transmission pairs among isolates that are either identical or closely related at the consensus genome level and is particularly effective at identifying isolates belonging to the same transmission chains. If appropriately employed, our approach has the potential to significantly reduce the number of potential transmission routes a contact tracer explores and can aid in early detection and interruption of transmission clusters if done in real-time. If this is done retrospectively, it can still help confirm or refute an outbreak, for example, two different transmission chains may have different implications for public health resources than one big one, particularly if the epidemiological characteristics are not the same and therefore different interventions are needed to 1) stop the current chains of transmission and 2) prevent future ones in those populations.

A unique component of this study was that it tracked the propagation of within-host variants over multiple consecutive transmissions until some became fixed SNVs or were eliminated by a population sweep. We observed that iSNVs were maintained over multiple transmission steps, which suggests that despite being small, the bottleneck size is greater than or comparable to the amount of within-host diversity in *Citrobacter rodentium*. This may explain why shared iSNVs were better at predicting whether isolates were from the same transmission chain or not. Although rare iSNVs are informative of transmission, using shared iSNVs means that data from a continuous scale (2.5-100%) are summarized into a binary (presence/absence) output and the presence of the iSNV in transmission chains obscures the transmission inference signal ^7,14^.

In outbreak investigations, there are three increasingly highly resolved levels of linkage between cases. At the most stringent we wish to be able to infer exact transmission pairs, yet it is also important and frequently valuable to tell the difference between transmission chains. Indeed, identifying whether two isolates are of the same outbreak are fundamental to the first principles of outbreak investigation. We note that the last of these is likely to be achievable with genomes in all cases other than those of very recent emergence (e.g., early pandemic). Our work shows that even among closely related isolates substantial further resolution is possible to the level of transmission chains and even individual transmission links. Our method was able to connect isolates that shared the same original infected host, which may contribute to the early identification of outbreaks. For example, the SARS-CoV2 pandemic has demonstrated how large numbers of downstream infections can arise from transmission clusters that emerge from super spreader events at mass gatherings ^28,29^. Early detection of transmission clusters can significantly curb downstream transmission, which will remain relevant as the pandemic progresses into endemicity. The next step in our work will be apply to it to datasets from infections in natural and/or human populations and incorporate temporal, clinical, and epidemiological data with our results to further discriminate true transmission pairs and clusters from spurious hits.

This work provides an advancement to track transmission chains. However, we recognize some limitations. Firstly, we only sampled a single snapshot just before each transmission. Therefore, we were unable to distinguish between diversity that arose due to a genetic bottleneck and genetic drift. This has implications for predicting what to expect to observe given a transmission event. Future work will include extending this approach to datasets where true transmission pairs and clusters are unknown. While the cost of deep sequencing remains relatively high and sequencing isn’t done routinely in many public health units, places with ongoing genomic surveillance can reserve this method for investigating suspected transmission clusters to guide rapid infection control responses. We further acknowledge that this method was only tested on one pathogen in a controlled laboratory experiment, and that results from clinical and field isolates may vary.

## Conclusion

We established multiple natural infection transmission chains and tracked the emergence and propagation of within-host variants until they became fixed SNVs or were eliminated by a population sweep. Beyond the presence and absence of within-host variants, we show that differences arising in the relative abundance of iSNVs can infer transmission clusters with high precision. Our results are an encouraging step towards higher precision contact tracing and early detection of transmission clusters. Our model can be incorporated into existing real-time sequencing frameworks and offer public health officials a quantifiable and actionable metric that can reliably infer transmission clusters.

## Methods

### Study design

Experiments were performed in accordance with the New Zealand Animal Welfare Act (1999) and institutional guidelines provided by the University of Auckland Animal Ethics Committee, which reviewed and approved these experiments under application R1003. We did not use any specific randomization process to allocate animals to a particular transmission chain or any specific strategies to minimize any confounding factors. All authors were also aware of the group allocation.

### Establishment of transmission chains

We have published a detailed description of our methods on the protocol repository website protocols.io ^30^, and a schematic of the experimental design is provided in Figure 1. Briefly, ten experimental transmission chains were established in female 6–7-week-old specific-pathogen-free (SPF) C57BL/6Elite mice (Vernon Jansen Unit, University of Auckland) using the bioluminescent *C. rodentium* strain ICC180. Initially, twelve seed mice were split into two groups and orally gavaged with *C. rodentium* (200 µL, ∼10^8^ colony forming units [CFU]) as previously described ^17^. One group was given drinking water supplemented with nalidixic acid (10 μg mL^−1^), which was refreshed every 2-3 days. Seven days post-gavage, the five animals with the highest ICC180 burden from each group (with nalidixic acid in the drinking water [N] or without [W]) were transferred to individual cages and designated transmission chain N1-5 or W1-5 and mouse (M)_1_/Effective Passage (EP)_1_. Each M_1_/EP_1_ animal was housed with an uninfected cage-mate, designated M_2_/EP_2_, for seven days to allow for transmission to occur via grooming and coprophagia. After seven days, the M_1_ animals were removed and humanely euthanized; each M_2_ animal was transferred to a clean cage and rehoused with an uninfected cage-mate, designated M_3_/EP_3._ We repeated this process until we reached M_22_/EP_22_. All animals in transmission chains N1-5 continued to receive drinking water supplemented with nalidixic acid, refreshed every 2-3 days.

### Monitoring of infection and transmission dynamics

We monitored mouse-to-mouse transmission of *C. rodentium* ICC180 by measuring luminescence and viable bacterial counts from stool samples recovered aseptically from individual animals, as previously described ^17^. Stool samples were also taken from infected animals on the day they were comingled with uninfected animals, suspended 1:1 in 50% glycerol and frozen at -80°C for genomic DNA extraction.

Where *C. rodentium* ICC180 failed to transmit between animals, we went back to the relevant frozen stool sample to produce an inoculum. For example, if *C. rodentium* failed to transmit during cohousing of animals M_3_/EP_3_ and M_4_/EP_4_, upon cohousing of M_4_/EP_4_ and M_5_/EP_5_, we orally gavaged animal M_5_ with an inoculum produced from the frozen stools of M_3_. Animal M_4_ was then redesignated M_4_/EP_NULL_ and M_5_ was redesignated M_5_/EP_4_ to account for the missed transmission step. Animals remained cohoused for humane reasons.

We also monitored transmission and infection dynamics using biphotonic imaging. Twice weekly we anaesthetized mice with gaseous isoflurane and measured bioluminescence using the IVIS® Kinetic imaging system (Perkin Elmer). Statistical analysis of infection and transmission dynamics data was carried out using the lme4 package (V 1.1-27.1) ^31^ in R (V 4.1). Linear mixed models were fitted to natural logarithms of bacterial load measures (CFU, bioluminescence), with fixed effects for antibiotic treatment, passage step, and time, and random effects for mouse and transmission chain. Analyses of transmission failure used logistic mixed models with a random effect for the chain.

### Whole-genome sequencing of C. rodentium ICC180 from infected animals

*C. rodentium* was obtained from frozen stool samples grown overnight at 37°C in 10mL LB-Lennox broth supplemented with kanamycin (50μg ml^−1^). Whole-genome DNA was extracted using a Qiagen DNeasy Blood and Tissue kit (Qiagen New Zealand Ltd, Auckland, New Zealand). The entire culture was used to capture diversity in shed stools as a surrogate for within-host diversity. Libraries were prepared using Nextera Flex (Illumina, San Diego, CA, USA). Library quality was checked by TapeStation and QPCR. Samples were sequenced at the Harvard University Bauer Core using the Illumina NovaSeq.

### Bioinformatic handling of sequence reads mapping and variant calling

We assessed reads quality using FASTQC. Trimmomatic (V 0.35) was employed to remove adapters and bases with a Phred quality score of < 33. Unpaired reads and sequences less than 50 bases long were discarded. The reads were mapped to the *C. rodentium* ICC168 reference genome (Accession Number: NC_013716.1) using bwa (V 0.7.17) with default parameters ^32^. Samtools (V 0.1.19)^33^ was used to filter unmapped reads. The GATK (V 4.0.2.1) toolkit preprocessing steps were applied to recalibrate base scores for mapped reads and perform joint variant calling ^34^ (Supplementary figure 4).

### Variant filtering and variant calling

We employed bcftools (version 1.9) to filter out variant sites with a QUAL score < 100, as well as sites with indels or multiple alternative alleles. Bedtools (v2.29.2)^35^ was used to mask variants within prophage regions of the reference genome, as identified by Magaziner *et al* (2019)^36^. Then, VCFtools (version 0.1.16) was used to remove consecutive variants within 100 bases window^37^. A custom Perl script was implemented to filter out sites that had fewer than 20 reads mapped. We identified 14 “Noisy” sites where their presence in multiple transmission chains and the allelic frequency at these sites fluctuated between 0.0 and 0.3 across successive pairs of samples (Supplementary figure 5). Consensus SNVs were called based on allelic frequency (reads mapping to the alternative allele) of at least 90%. iSNVs were designated as *loci* where the allelic frequency ranged from 2.5% to 90% of reads mapping to the alternative allele. We generated an SNV alignment based on the consensus call for each genome.

### Phylogenetic tree reconstruction

The core SNPs were aligned with MAFFT (V 7.467). Alignment was used as input to RAxML (V 8.2.12) to reconstruct the phylogenetic tree using the general time-reversible model and gamma correction ^38,39^. Since we used only variable sites as input, we used ASC_GTRGAMMA to correct ascertainment bias with the Paul Lewis correction. The isolate ICC180 was used as the outgroup. One thousand bootstrap replicates were generated to assess the significance of internal nodes.

### Pairwise comparisons and bottleneck estimates

Statistical analysis of whole genome sequencing data was carried out in R (V 4.2.1). A pairwise SNV distance matrix based on presence and absence was calculated from all samples using Mega7 (V 7.0.267) ^40^. For each pairwise comparison, we computed additional metrics to enhance our ability to distinguish transmissions pairs from non-transmission pairs. First, each transmission step was taken as a single unit in time, and the sum of transmission steps separating the two isolates was recorded. For isolates in separate transmission chains, this was the sum of cumulative transmission steps from the index strain up to the isolate being queried in both chains. Then, we computed the number of shared iSNVs and variable sites as the number of sites where both strains had an iSNV and SNV. We defined allelic frequency as the proportion of reads mapping to the alternative allele. At each variable locus, we computed the change in the allelic frequency between the two isolates. We then calculated the mean change in allelic ratio (θ), based on the number of loci where the isolates had different allelic frequencies. Finally, we employed the beta-binomial method to infer bottleneck sizes for transmission pairs, based on allelic frequencies in the donor and recipient (https://github.com/weissmanlab/BB_bottleneck)^15^.

### Bayesian framework to infer the likelihood of transmission

We used the mean change allelic frequency to infer the posterior probability of transmission (which we refer to as likelihood of transmission). Let *P(T)* be the proportion of comparisons that are transmission pairs, P(θ) be the probability of observing θ and P(θ|Τ) the probability of observing θ in a transmission pair. The prior probability density distributions P(θ) and P(θ|Τ) were inferred by fitting distribution of θ for non-transmission and transmission to a truncated normal distribution using the fitdist package in R. We then calculated the posterior probability P(T|θ) of observing a transmission given the mean change in allelic frequency using Bayes theorem (Equation 1).

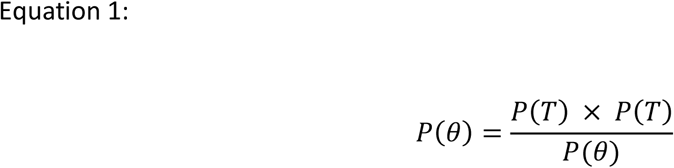

Similarly, the Bayes theorem was also used to infer P(T|V), where V represented the number of shared sites that were either an iSNV or a SNV in both isolates (Equation 2):

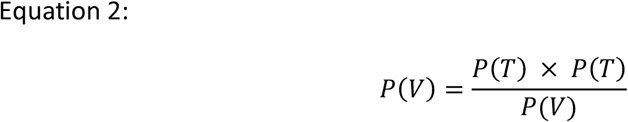

The obtained likelihood P(Τ|θ) was used to reconstruct the transmission network (Figure 5B). At this step, we only considered transmissions that were within potential donors within three steps. We then ranked all comparisons based on transmission likelihood. The best potential donor was the one with the highest transmission likelihood. We only allowed donors in transmission steps lower than recipients.

## Funding

This work was supported by an R01 grant from the National Institutes of Health, awarded to WPH (R01AI128344), and grants-in-aid from the University of Auckland’s Faculty of Medical and Health Sciences (9802-3701152) and the Maurice Wilkins Centre for Molecular Biodiscovery (9431-48516), awarded to SW.

## Acknowledgements

The authors acknowledge the staff of the Vernon Jenson Unit for breeding the mice required for this study, and for the care and compassion they show to all the animals in the Unit. The authors also acknowledge the 222 mice used in this research, without whom this work wouldn’t have been possible.

## Data availability

The raw infection data from this study have been uploaded to Figshare (https://doi.org/10.17608/k6.auckland.21350790.v1). The raw data sequencing reads from the study have been uploaded to the NCBI short read archives under the Bioproject ID: PRJNA884719.

## Supplementary figures

**Supplementary Figure 1.**
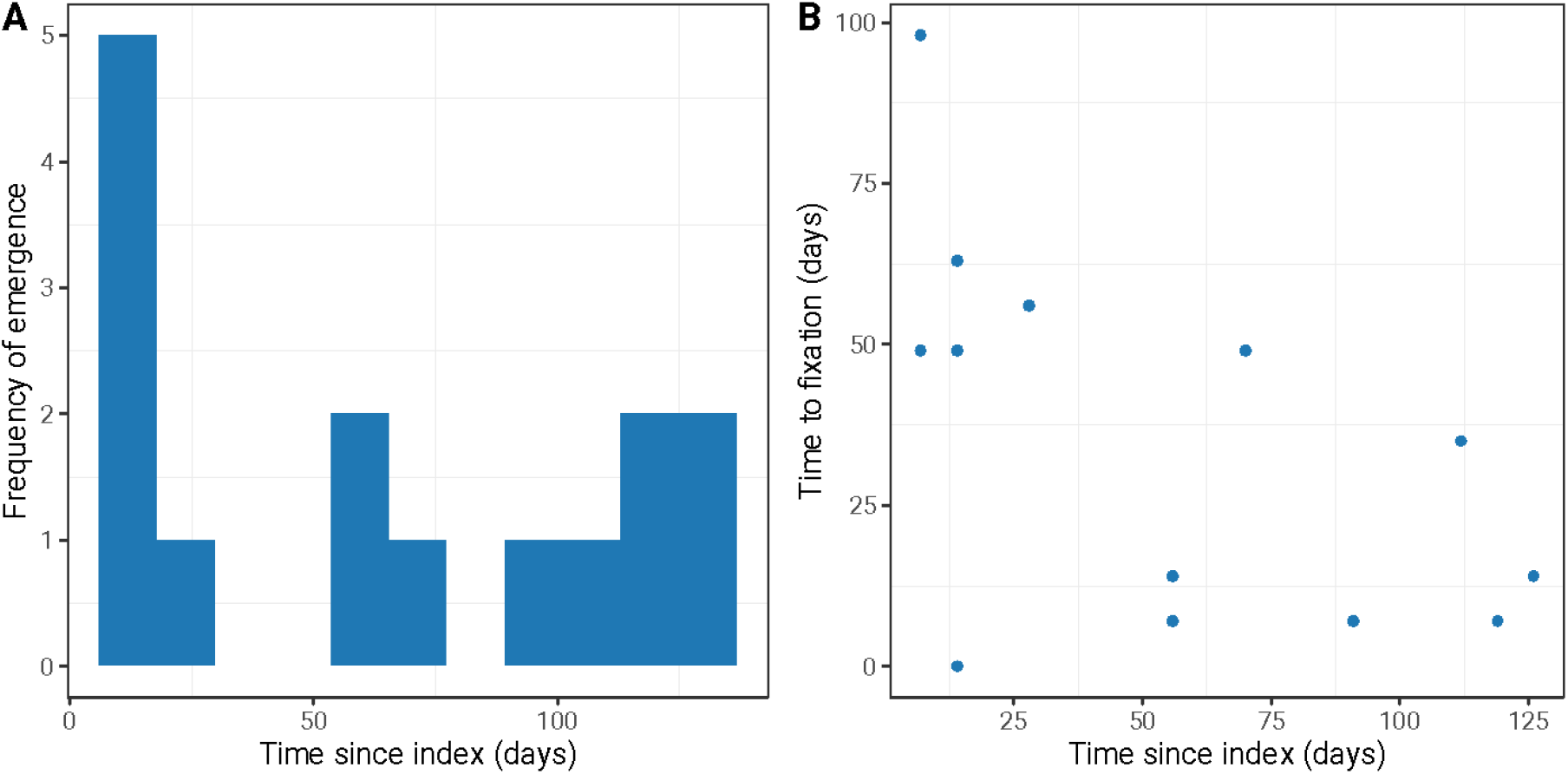
Emergence of iSNVs and the time it takes to become fixed SNVs. A) Histogram illustrating the frequency with which iSNVs were emerging at a time t (days) since the index. B) Scatter plot of when iSNVs emerge in the transmission chain and how long it takes them to reach fixation.

**Supplementary Figure 2.**
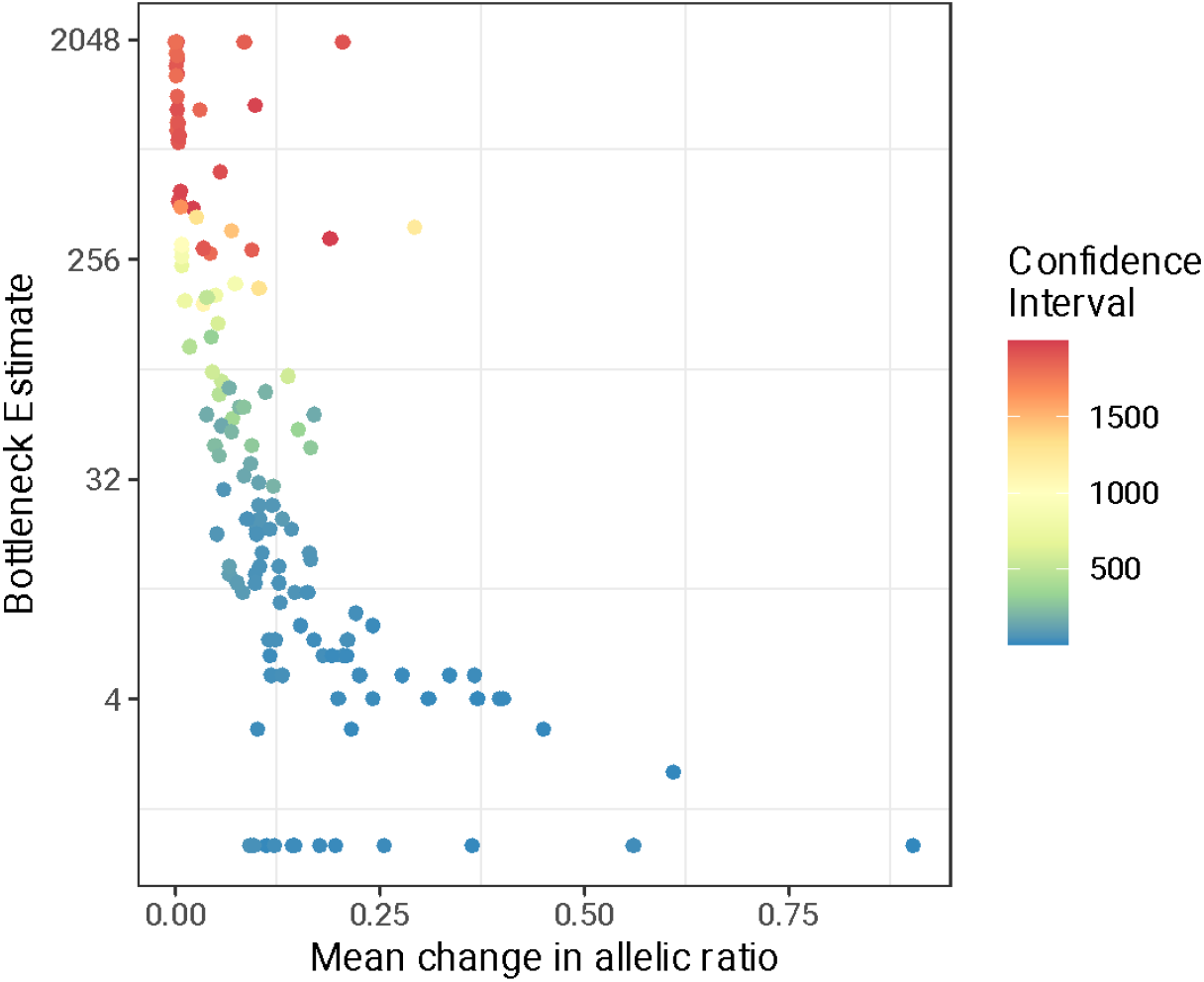
Genetic bottleneck estimates for transmission pairs inferred from allelic ratio colored by the width of the confidence intervals.

**Supplementary Figure 3.**
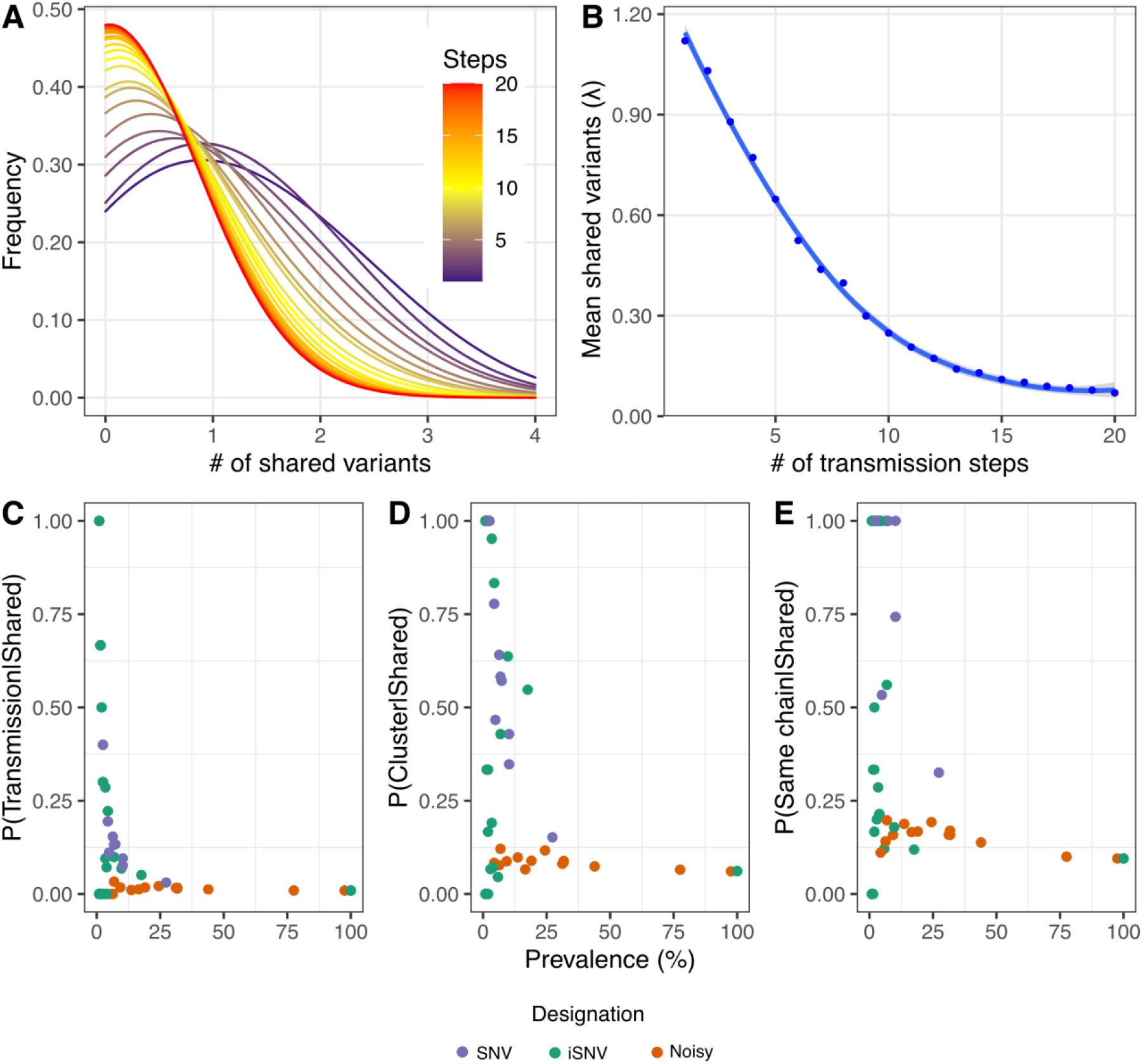
Counting shared genomic variants to infer transmission and the change in the number of shared variants over multiple transmission steps. A) Density plot showing the distribution of shared genomic variants, grouped by the number of transmission steps linking pairs. B) The Poisson fitted the mean number of shared variants for pairs linked by the number of transmission steps. C-E) The probability that two isolates are of a transmission pair, from the same transmission cluster or the same transmission chain, given the presence of a variant in both isolates, at the given loci. The x-axis shows the prevalence of variants at the *loci* across the entire dataset.

**Supplementary Table S1.** Transmission and Infection dynamics by individual animal Key: OG, oral gavage; NT, natural transmission; NI, not infected; NA, not applicable.

| Transmission chain | Animal | Effective Passage (EP) | Weight (g) (Median [range]) | Weight change since start (%) | Weight change at peak (%) | Route of infection | Viable counts prior to co-mingling (CFU g<sup>-1</sup>) |
| --- | --- | --- | --- | --- | --- | --- | --- |
| N1 | M<sub>1</sub> | EP<sub>1</sub> | 18.30 (1.20) | 4.95 | 0.55 | OG | 2.67 x 10<sup>9</sup> |
|  | M<sub>2</sub> | EP<sub>2</sub> | 16.00 (0.50) | -1.88 | -1.88 | NT | 1.07 x 10<sup>9</sup> |
|  | M<sub>3</sub> | EP<sub>3</sub> | 19.20 (1.00) | 1.52 | -2.54 | NT | 5.33 x 10<sup>8</sup> |
|  | M<sub>4</sub> | EP<sub>4</sub> | 17.35 (1.70) | 6.94 | 0.00 | NT | 1.67 x 10<sup>9</sup> |
|  | M<sub>5</sub> | EP<sub>5</sub> | 18.10 (2.30) | 3.78 | -0.54 | NT | 1.00 x 10<sup>9</sup> |
|  | M<sub>6</sub> | EP<sub>6</sub> | 15.50 (0.70) | 2.56 | -1.92 | NT | 1.87 x 10<sup>9</sup> |
|  | M<sub>7</sub> | EP<sub>7</sub> | 18.55 (1.40) | -5.70 | -2.59 | NT | 1.47 x 10<sup>9</sup> |
|  | M<sub>8</sub> | EP<sub>8</sub> | 18.55 (1.40) | -1.56 | -5.21 | NT | 9.50 x 10<sup>8</sup> |
|  | M<sub>9</sub> | EP<sub>9</sub> | 17.65 (0.60) | -0.56 | -1.69 | NT | 8.17 x 10<sup>8</sup> |
|  | M<sub>10</sub> | EP<sub>10</sub> | 17.95 (1.00) | 0.00 | 0.00 | NT | 5.50 x 10<sup>8</sup> |
|  | M<sub>11</sub> | EP<sub>11</sub> | 17.05 (0.70) | 1.18 | 2.94 | NT | 7.00 x 10<sup>8</sup> |
|  | M<sub>12</sub> | EP<sub>12</sub> | 17.20 (1.00) | 2.87 | -1.15 | NT | 1.23 x 10<sup>8</sup> |
|  | M<sub>13</sub> | EP<sub>13</sub> | 14.75 (1.30) | 3.40 | 2.72 | NT | 2.35 x 10<sup>9</sup> |
|  | M<sub>14</sub> | EP<sub>14</sub> | 17.00 (1.70) | 5.29 | 1.76 | NT | 2.08 x 10<sup>9</sup> |
|  | M<sub>15</sub> | EP<sub>15</sub> | 18.40 (1.90) | 2.67 | -7.49 | NT | 1.20 x 10<sup>9</sup> |
|  | M<sub>16</sub> | EP<sub>16</sub> | 18.90 (0.90) | 1.03 | -2.58 | NT | 1.63 x 10<sup>9</sup> |
|  | M<sub>17</sub> | EP<sub>17</sub> | 18.05 (0.80) | 1.68 | -1.68 | NT | 1.47 x 10<sup>9</sup> |
|  | M<sub>18</sub> | EP<sub>18</sub> | 19.25 (0.70) | 2.62 | -1.05 | NT | 1.30 x 10<sup>9</sup> |
|  | M<sub>19</sub> | EP<sub>19</sub> | 18.15 (1.50) | -3.87 | 0.55 | NT | 4.00 x 10<sup>8</sup> |
|  | M<sub>20</sub> | EP<sub>20</sub> | 17.80 (0.60) | 1.71 | 1.71 | NT | 5.83 x 10<sup>7</sup> |
|  | M<sub>21</sub> | EP<sub>21</sub> | 17.25 (1.20) | 7.06 | 5.29 | NT | 1.10 x 10<sup>9</sup> |
|  | M<sub>22</sub> | EP<sub>22</sub> | 17.15 (0.90) | -1.88 | 3.49 | NT | NA |
| N2 | M<sub>1</sub> | EP<sub>1</sub> | 19.70 (2.20) | 3.17 | 4.76 | OG | 1.35 x 10<sup>9</sup> |
|  | M<sub>2</sub> | EP<sub>2</sub> | 16.40 (2.40) | 4.22 | -4.22 | NT | 5.00 x 10<sup>8</sup> |
|  | M<sub>3</sub> | EP<sub>3</sub> | 19.60 (0.90) | 2.55 | -2.04 | NT | 7.00 x 10<sup>8</sup> |
|  | M<sub>4</sub> | EP<sub>4</sub> | 18.20 (0.30) | -0.55 | 1.10 | NT | 1.45 x 10<sup>9</sup> |
|  | **M<sub>5</sub>** | **EP<sub>NULL</sub>** | **19.55 (0.90)** | **3.14** | 0.52 | **NI** | **3.33 x 10<sup>2</sup>** |
|  | M<sub>6</sub> | EP<sub>5</sub> | 16.60 (1.60) | 6.63 | -3.01 | OG | 1.15 x 10<sup>9</sup> |
|  | M<sub>7</sub> | EP<sub>6</sub> | 20.75 (1.10) | 1.91 | -0.96 | NT | 2.28 x 10<sup>9</sup> |
|  | M<sub>8</sub> | EP<sub>7</sub> | 17.40 (0.70) | 1.15 | 0.57 | NT | 8.67 x 10<sup>8</sup> |
|  | M<sub>9</sub> | EP<sub>8</sub> | 17.20 (2.00) | 8.67 | -2.89 | NT | 2.93 x 10<sup>9</sup> |
|  | M<sub>10</sub> | EP<sub>9</sub> | 17.40 (1.10) | -2.26 | -1.13 | NT | 6.67 x 10<sup>8</sup> |
|  | M<sub>11</sub> | EP<sub>10</sub> | 16.60 (1.20) | -0.59 | -7.06 | NT | 1.05 x 10<sup>9</sup> |
|  | M<sub>12</sub> | EP<sub>11</sub> | 18.35 (1.40) | 4.89 | -0.54 | NT | 4.50 x 10<sup>8</sup> |
|  | M<sub>13</sub> | EP<sub>12</sub> | 19.55 (1.40) | 7.53 | 0.54 | NT | 6.67 x 10<sup>8</sup> |
|  | M<sub>14</sub> | EP<sub>13</sub> | 18.55 (0.70) | -0.54 | 0.54 | NT | 1.32 x 10<sup>9</sup> |
|  | M<sub>15</sub> | EP<sub>14</sub> | 19.15 (1.10) | 2.59 | -3.11 | NT | 2.83 x 10<sup>8</sup> |
|  | M<sub>16</sub> | EP<sub>15</sub> | 18.05 (0.30) | 0.55 | -1.10 | NT | 7.00 x 10<sup>7</sup> |
|  | M<sub>17</sub> | EP<sub>16</sub> | 17.30 (0.90) | 2.89 | 0.58 | NT | 1.47 x 10<sup>9</sup> |
|  | M<sub>18</sub> | EP<sub>17</sub> | 19.00 (1.20) | 4.35 | 4.35 | NT | 9.00 x 10<sup>6</sup> |
|  | M<sub>19</sub> | EP<sub>18</sub> | 17.70 (0.80) | 0.56 | 0.00 | NT | 4.33 x 10<sup>8</sup> |
|  | M<sub>20</sub> | EP<sub>19</sub> | 18.10 (1.10) | 5.65 | -0.56 | NT | 4.83 x 10<sup>8</sup> |
|  | M<sub>21</sub> | EP<sub>20</sub> | 18.55 (1.00) | 2.15 | -3.23 | NT | 4.67 x 10<sup>8</sup> |
|  | M<sub>22</sub> | EP<sub>21</sub> | 19.50 (1.10) | -5.53 | -5.53 | NT | NA |
| N3 | M<sub>1</sub> | EP<sub>1</sub> | 18.30 (1.60) | 5.98 | 0.54 | OG | 4.00 x 10<sup>9</sup> |
|  | M<sub>2</sub> | EP<sub>2</sub> | 16.60 (0.60) | 3.68 | 3.68 | NT | 4.33 x 10<sup>8</sup> |
|  | M<sub>3</sub> | EP<sub>3</sub> | 18.40 (0.70) | 0.53 | -2.67 | NT | 9.00 x 10<sup>7</sup> |
|  | M<sub>4</sub> | EP<sub>4</sub> | 15.80 (0.80) | 3.18 | 2.55 | NT | 7.33 x 10<sup>7</sup> |
|  | M<sub>5</sub> | EP<sub>5</sub> | 19.20 (0.90) | 2.07 | -0.52 | NT | 1.83 x 10<sup>9</sup> |
|  | M<sub>6</sub> | EP<sub>6</sub> | 15.75 (1.50) | 9.62 | 0.00 | NT | 9.83 x 10<sup>8</sup> |
|  | M<sub>7</sub> | EP<sub>7</sub> | 19.05 (1.20) | 1.55 | -1.55 | NT | 1.13 x 10<sup>9</sup> |
|  | M<sub>8</sub> | EP<sub>8</sub> | 16.70 (1.50) | 7.83 | 1.20 | NT | 9.33 x 10<sup>8</sup> |
|  | M<sub>9</sub> | EP<sub>9</sub> | 19.00 (1.20) | 5.79 | -0.53 | NT | 8.17 x 10<sup>8</sup> |
|  | M<sub>10</sub> | EP<sub>10</sub> | 17.90 (0.90) | -1.10 | 0.55 | NT | 6.67 x 10<sup>8</sup> |
|  | M<sub>11</sub> | EP<sub>11</sub> | 17.95 (1.20) | 5.59 | 3.35 | NT | 1.23 x 10<sup>8</sup> |
|  | M<sub>12</sub> | EP<sub>12</sub> | 18.90 (1.30) | 4.76 | -2.12 | NT | 9.67 x 10<sup>8</sup> |
|  | M<sub>13</sub> | EP<sub>13</sub> | 13.30 (1.30) | -5.76 | 0.00 | NT | 6.83 x 10<sup>8</sup> |
|  | M<sub>14</sub> | EP<sub>14</sub> | 18.55 (1.00) | -0.54 | -2.69 | NT | 4.00 x 10<sup>8</sup> |
|  | M<sub>15</sub> | EP<sub>15</sub> | 19.80 (1.30) | 1.48 | -4.93 | NT | 4.00 x 10<sup>8</sup> |
|  | M<sub>16</sub> | EP<sub>16</sub> | 18.20 (1.10) | 4.42 | -1.66 | NT | 4.00 x 10<sup>8</sup> |
|  | M<sub>17</sub> | EP<sub>17</sub> | 18.70 (1.00) | 3.74 | -1.60 | NT | 6.33 x 10<sup>7</sup> |
|  | M<sub>18</sub> | EP<sub>18</sub> | 18.80 (1.40) | -0.51 | -2.05 | NT | 2.55 x 10<sup>8</sup> |
|  | M<sub>19</sub> | EP<sub>19</sub> | 20.10 (0.60) | 1.49 | 1.49 | NT | 9.17 x 10<sup>7</sup> |
|  | **M<sub>20</sub>** | **EP<sub>NULL</sub>** | **18.40 (1.80)** | **6.67** | **1.11** | **NI** | **1.00 x 10<sup>3</sup>** |
|  | M<sub>21</sub> | EP<sub>20</sub> | 18.50 (1.50) | 6.56 | 1.09 | OG | 5.83 x 10<sup>8</sup> |
|  | M<sub>22</sub> | EP<sub>21</sub> | 17.30 (1.00) | 2.86 | 2.86 | NT | NA |
| N4 | M<sub>1</sub> | EP<sub>1</sub> | 18.50 (2.50) | -2.70 | -5.41 | OG | 2.50 x 10<sup>9</sup> |
|  | M<sub>2</sub> | EP<sub>2</sub> | 18.10 (1.30) | 7.34 | 2.26 | NT | 6.00 x 10<sup>8</sup> |
|  | M<sub>3</sub> | EP<sub>3</sub> | 18.90 (1.10) | 3.68 | -2.11 | NT | 7.00 x 10<sup>8</sup> |
|  | M<sub>4</sub> | EP<sub>4</sub> | 16.75 (1.00) | 6.02 | 1.20 | NT | 8.33 x 10<sup>8</sup> |
|  | M<sub>5</sub> | EP<sub>5</sub> | 19.30 (1.00) | -1.02 | -2.54 | NT | 1.42 x 10<sup>9</sup> |
|  | M<sub>6</sub> | EP<sub>6</sub> | 16.90 (1.30) | 4.14 | -3.55 | NT | 1.63 x 10<sup>9</sup> |
|  | M<sub>7</sub> | EP<sub>7</sub> | 19.60 (0.90) | 3.09 | 1.55 | NT | 3.17 x 10<sup>8</sup> |
|  | M<sub>8</sub> | EP<sub>8</sub> | 16.00 (0.90) | 3.75 | 0.63 | NT | 4.50 x 10<sup>7</sup> |
|  | M<sub>9</sub> | EP<sub>9</sub> | 19.05 (2.10) | 4.71 | 2.62 | NT | 2.32 x 10<sup>9</sup> |
|  | M<sub>10</sub> | EP<sub>10</sub> | 16.85 (1.20) | 2.91 | -2.33 | NT | 7.67 x 10<sup>8</sup> |
|  | M<sub>11</sub> | EP<sub>11</sub> | 16.75 (1.60) | -6.78 | -3.95 | NT | 1.02 x 10<sup>9</sup> |
|  | M<sub>12</sub> | EP<sub>12</sub> | 15.80 (1.70) | -8.28 | -4.73 | NT | 1.58 x 10<sup>7</sup> |
|  | M<sub>13</sub> | EP<sub>13</sub> | 18.70 (0.90) | -1.60 | 0.53 | NT | 6.83 x 10<sup>8</sup> |
|  | M<sub>14</sub> | EP<sub>14</sub> | 18.00 (1.50) | 7.30 | 5.06 | NT | 3.00 x 10<sup>9</sup> |
|  | M<sub>15</sub> | EP<sub>15</sub> | 16.05 (0.90) | 4.38 | -1.25 | NT | 8.33 x 10<sup>7</sup> |
|  | M<sub>16</sub> | EP<sub>16</sub> | 17.70 (1.00) | 5.78 | 5.78 | NT | 1.00 x 10<sup>9</sup> |
|  | M<sub>17</sub> | EP<sub>17</sub> | 17.45 (0.70) | 0.57 | -0.57 | NT | 1.10 x 10<sup>9</sup> |
|  | M<sub>18</sub> | EP<sub>18</sub> | 17.25 (1.40) | 7.06 | 1.76 | NT | 1.63 x 10<sup>9</sup> |
|  | M<sub>19</sub> | EP<sub>19</sub> | 18.10 (1.30) | 3.83 | -3.28 | NT | 1.08 x 10<sup>8</sup> |
|  | M<sub>20</sub> | EP<sub>20</sub> | 19.70 (1.20) | 3.05 | 0.00 | NT | 6.00 x 10<sup>7</sup> |
|  | M<sub>21</sub> | EP<sub>21</sub> | 16.90 (2.10) | 12.12 | -0.61 | NT | 1.47 x 10<sup>8</sup> |
|  | M<sub>22</sub> | EP<sub>22</sub> | 17.65 (0.10) | 0.57 | 0.57 | NT | NA |
| N5 | M<sub>1</sub> | EP<sub>1</sub> | 18.80 (2.60) | -1.03 | -13.40 | OG | 2.50 x 10<sup>9</sup> |
|  | M<sub>2</sub> | EP<sub>2</sub> | 18.40 (1.50) | 2.73 | 3.28 | NT | 2.00 x 10<sup>8</sup> |
|  | M<sub>3</sub> | EP<sub>3</sub> | 17.90 (0.60) | 1.68 | -1.68 | NT | 2.67 x 10<sup>8</sup> |
|  | M<sub>4</sub> | EP<sub>4</sub> | 18.30 (0.60) | 1.64 | -0.55 | NT | 2.33 x 10<sup>8</sup> |
|  | M<sub>5</sub> | EP<sub>5</sub> | 17.90 (1.20) | -0.54 | -4.84 | NT | 1.53 x 10<sup>9</sup> |
|  | M<sub>6</sub> | EP<sub>6</sub> | 17.00 (1.50) | 7.10 | 0.00 | NT | 5.83 x 10<sup>8</sup> |
|  | **M<sub>7</sub>** | **EP<sub>NULL</sub>** | **16.85 (0.80)** | **1.79** | **-0.60** | **NI** | **<1.67 x 10<sup>2</sup>** |
|  | M<sub>8</sub> | EP<sub>7</sub> | 19.05 (1.30) | 4.23 | 4.76 | OG | 2.07 x 10<sup>9</sup> |
|  | M<sub>9</sub> | EP<sub>8</sub> | 19.45 (1.10) | 4.62 | 2.05 | NT | 4.50 x 10<sup>8</sup> |
|  | M<sub>10</sub> | EP<sub>9</sub> | 17.90 (1.90) | 10.11 | 3.37 | NT | 5.33 x 10<sup>8</sup> |
|  | M<sub>11</sub> | EP<sub>10</sub> | 17.75 (1.30) | 7.56 | 4.65 | NT | 5.17 x 10<sup>8</sup> |
|  | M<sub>12</sub> | EP<sub>11</sub> | 18.60 (1.80) | 3.17 | -5.29 | NT | 4.17 x 10<sup>8</sup> |
|  | M<sub>13</sub> | EP<sub>12</sub> | 18.30 (1.40) | 7.95 | 5.68 | NT | 7.33 x 10<sup>8</sup> |
|  | M<sub>14</sub> | EP<sub>13</sub> | 18.20 (1.30) | 0.00 | -6.99 | NT | 1.82 x 10<sup>8</sup> |
|  | **M<sub>15</sub>** | **EP<sub>NULL</sub>** | **17.90 (1.00)** | **4.57** | **0.57** | **NI** | **6.67 x 10<sup>2</sup>** |
|  | M<sub>16</sub> | EP<sub>14</sub> | 18.35 (1.20) | 4.92 | 0.55 | OG | 2.40 x 10<sup>9</sup> |
|  | M<sub>17</sub> | EP<sub>15</sub> | 17.35 (0.70) | 2.31 | 1.16 | NT | 4.17 x 10<sup>8</sup> |
|  | M<sub>18</sub> | EP<sub>16</sub> | 19.15 (2.10) | 11.35 | 2.70 | NT | 1.62 x 10<sup>8</sup> |
|  | M<sub>19</sub> | EP<sub>17</sub> | 19.55 (1.10) | 0.51 | 1.53 | NT | 8.17 x 10<sup>7</sup> |
|  | M<sub>20</sub> | EP<sub>18</sub> | 18.40 (0.70) | 1.64 | -2.19 | NT | 4.17 x 10<sup>8</sup> |
|  | M<sub>21</sub> | EP<sub>19</sub> | 18.55 (1.60) | 8.84 | 2.21 | NT | 2.50 x 10<sup>8</sup> |
|  | M<sub>22</sub> | EP<sub>20</sub> | 19.20 (0.20) | 1.05 | 1.05 | NT | NA |
| W1 | M<sub>1</sub> | EP<sub>1</sub> | 17.10 (2.20) | 2.87 | -5.17 | OG | 1.03 x 10<sup>9</sup> |
|  | M<sub>2</sub> | EP<sub>2</sub> | 17.80 (1.10) | 6.36 | 4.05 | NT | 9.00 x 10<sup>7</sup> |
|  | M<sub>3</sub> | EP<sub>3</sub> | 17.70 (1.00) | 1.69 | -1.12 | NT | 5.50 x 10<sup>8</sup> |
|  | M<sub>4</sub> | EP<sub>4</sub> | 18.90 (0.30) | 0.00 | -1.59 | NT | 5.17 x 10<sup>8</sup> |
|  | M<sub>5</sub> | EP<sub>5</sub> | 17.60 (1.13) | 2.23 | 0.00 | NT | 2.33 x 10<sup>9</sup> |
|  | M<sub>6</sub> | EP<sub>6</sub> | 18.15 (1.00) | 5.03 | 1.68 | NT | 1.07 x 10<sup>9</sup> |
|  | M<sub>7</sub> | EP<sub>7</sub> | 20.15 (0.90) | 1.47 | -2.94 | NT | 1.22 x 10<sup>9</sup> |
|  | M<sub>8</sub> | EP<sub>8</sub> | 18.60 (1.40) | 6.04 | -1.65 | NT | 5.83 x 10<sup>8</sup> |
|  | M<sub>9</sub> | EP<sub>9</sub> | 18.15 (1.10) | 2.73 | 2.19 | NT | 6.00 x 10<sup>8</sup> |
|  | M<sub>10</sub> | EP<sub>10</sub> | 18.50 (1.00) | 4.32 | 0.54 | NT | 2.50 x 10<sup>8</sup> |
|  | M<sub>11</sub> | EP<sub>11</sub> | 16.50 (1.00) | 6.25 | 1.25 | NT | 6.67 x 10<sup>8</sup> |
|  | M<sub>12</sub> | EP<sub>12</sub> | 18.80 (1.70) | 6.91 | 0.00 | NT | 3.83 x 10<sup>8</sup> |
|  | M<sub>13</sub> | EP<sub>13</sub> | 17.15 (1.10) | 2.92 | -3.51 | NT | 2.38 x 10<sup>9</sup> |
|  | M<sub>14</sub> | EP<sub>14</sub> | 17.80 (1.10) | 0.56 | -1.12 | NT | 9.83 x 10<sup>7</sup> |
|  | M<sub>15</sub> | EP<sub>15</sub> | 16.85 (2.30) | 10.18 | 1.80 | NT | 7.67 x 10<sup>8</sup> |
|  | M<sub>16</sub> | EP<sub>16</sub> | 16.85 (0.70) | 2.37 | 0.59 | NT | 2.33 x 10<sup>8</sup> |
|  | M<sub>17</sub> | EP<sub>17</sub> | 18.00 (0.70) | -1.64 | -2.19 | NT | 8.50 x 10<sup>8</sup> |
|  | M<sub>18</sub> | EP<sub>18</sub> | 17.70 (1.10) | 2.23 | -3.91 | NT | 8.67 x 10<sup>8</sup> |
|  | M<sub>19</sub> | EP<sub>19</sub> | 19.05 (0.60) | 1.05 | -0.53 | NT | 5.50 x 10<sup>8</sup> |
|  | M<sub>20</sub> | EP<sub>20</sub> | 16.90 (1.50) | 3.05 | 3.66 | NT | 8.17 x 10<sup>8</sup> |
|  | M<sub>21</sub> | EP<sub>21</sub> | 18.40 (1.60) | 3.83 | -3.28 | NT | 6.50 x 10<sup>8</sup> |
|  | M<sub>22</sub> | EP<sub>22</sub> | 19.25 (0.60) | 2.60 | 2.60 | NT | NA |
| W2 | M<sub>1</sub> | EP<sub>1</sub> | 19.20 (0.50) | 1.56 | 1.04 | OG | 5.33 x 10<sup>9</sup> |
|  | M<sub>2</sub> | EP<sub>2</sub> | 18.10 (0.80) | 3.89 | 3.89 | NT | 4.17 x 10<sup>8</sup> |
|  | M<sub>3</sub> | EP<sub>3</sub> | 17.10 (1.40) | 5.85 | 1.75 | NT | 2.00 x 10<sup>8</sup> |
|  | M<sub>4</sub> | EP<sub>4</sub> | 18.55 (1.20) | 5.46 | 3.28 | NT | 8.33 x 10<sup>8</sup> |
|  | M<sub>5</sub> | EP<sub>5</sub> | 17.90 (1.10) | 1.69 | -0.56 | NT | 6.50 x 10<sup>8</sup> |
|  | **M<sub>6</sub>** | **EP<sub>NULL</sub>** | **16.85 (0.70)** | **1.78** | **-1.78** | **NI** | **<1.67 x 10<sup>2</sup>** |
|  | M<sub>7</sub> | EP<sub>6</sub> | 17.95 (0.90) | -1.10 | -2.20 | OG | 3.83 x 10<sup>8</sup> |
|  | M<sub>8</sub> | EP<sub>7</sub> | 15.85 (2.20) | 12.10 | -1.91 | NT | 3.17 x 10<sup>8</sup> |
|  | M<sub>9</sub> | EP<sub>8</sub> | 15.60 (1.40) | 5.13 | -0.64 | NT | 6.00 x 10<sup>8</sup> |
|  | M<sub>10</sub> | EP<sub>9</sub> | 17.15 (1.20) | 4.09 | 2.34 | NT | 8.17 x 10<sup>8</sup> |
|  | M<sub>11</sub> | EP<sub>10</sub> | 17.15 (0.90) | 0.00 | -1.16 | NT | 1.08 x 10<sup>9</sup> |
|  | M<sub>12</sub> | EP<sub>11</sub> | 17.50 (1.10) | 5.14 | 1.14 | NT | 2.83 x 10<sup>8</sup> |
|  | M<sub>13</sub> | EP<sub>12</sub> | 18.15 (0.80) | 1.63 | -2.72 | NT | 1.22 x 10<sup>9</sup> |
|  | M<sub>14</sub> | EP<sub>13</sub> | 20.55 (0.90) | 3.92 | 1.96 | NT | 1.12 x 10<sup>9</sup> |
|  | M<sub>15</sub> | EP<sub>14</sub> | 18.25 (1.60) | 8.24 | 0.00 | NT | 7.50 x 10<sup>8</sup> |
|  | M<sub>16</sub> | EP<sub>15</sub> | 17.95 (1.10) | 6.25 | 3.41 | NT | 5.83 x 10<sup>8</sup> |
|  | M<sub>17</sub> | EP<sub>16</sub> | 19.20 (1.30) | 2.06 | -2.06 | NT | 1.75 x 10<sup>9</sup> |
|  | M<sub>18</sub> | EP<sub>17</sub> | 16.50 (0.90) | -0.60 | -1.80 | NT | 1.38 x 10<sup>9</sup> |
|  | M<sub>19</sub> | EP<sub>18</sub> | 18.30 (1.10) | 4.37 | 0.00 | NT | 3.17 x 10<sup>8</sup> |
|  | M<sub>20</sub> | EP<sub>19</sub> | 18.90 (0.50) | 1.60 | 2.13 | NT | 5.33 x 10<sup>8</sup> |
|  | M<sub>21</sub> | EP<sub>20</sub> | 18.10 (1.50) | 6.63 | 0.55 | NT | 9.83 x 10<sup>8</sup> |
|  | M<sub>22</sub> | EP<sub>21</sub> | 19.60 (0.40) | -1.53 | -1.53 | NT | NA |
| W3 | M<sub>1</sub> | EP<sub>1</sub> | 18.70 (0.70) | 1.60 | 0.53 | OG | 2.50 x 10<sup>9</sup> |
|  | M<sub>2</sub> | EP<sub>2</sub> | 18.80 (0.80) | 2.73 | 3.83 | NT | 1.00 x 10<sup>8</sup> |
|  | **M<sub>3</sub>** | **EP<sub>NULL</sub>** | **17.80 (1.10)** | **1.70** | **0.00** | **NI** | **<1.67 x 10<sup>2</sup>** |
|  | M<sub>4</sub> | EP<sub>3</sub> | 16.90 (0.80) | 0.59 | -1.78 | OG | 1.50 x 10<sup>8</sup> |
|  | M<sub>5</sub> | EP<sub>4</sub> | 18.15 (1.40) | 7.22 | 0.00 | NT | 1.10 x 10<sup>9</sup> |
|  | M<sub>6</sub> | EP<sub>5</sub> | 17.00 (1.20) | 4.07 | -2.33 | NT | 1.08 x 10<sup>9</sup> |
|  | **M<sub>7</sub>** | **EP<sub>NULL</sub>** | **18.60 (1.10)** | **2.66** | **-1.60** | **NI** | **<1.67 x 10<sup>2</sup>** |
|  | M<sub>8</sub> | EP<sub>6</sub> | 18.30 (1.70) | 5.49 | -1.65 | OG | 2.00 x 10<sup>9</sup> |
|  | M<sub>9</sub> | EP<sub>7</sub> | 18.35 (1.00) | 4.92 | 0.55 | NT | 7.00 x 10<sup>8</sup> |
|  | M<sub>10</sub> | EP<sub>8</sub> | 16.25 (1.20) | 4.27 | -0.61 | NT | 1.67 x 10<sup>8</sup> |
|  | M<sub>11</sub> | EP<sub>9</sub> | 18.75 (0.80) | 2.69 | -0.54 | NT | 1.08 x 10<sup>8</sup> |
|  | M<sub>12</sub> | EP<sub>10</sub> | 17.70 (1.00) | 2.79 | -1.12 | NT | 7.67 x 10<sup>8</sup> |
|  | M<sub>13</sub> | EP<sub>11</sub> | 16.50 (0.80) | 2.47 | -1.23 | NT | 2.15 x 10<sup>9</sup> |
|  | M<sub>14</sub> | EP<sub>12</sub> | 17.00 (1.50) | 2.94 | 0.00 | NT | 7.67 x 10<sup>8</sup> |
|  | M<sub>15</sub> | EP<sub>13</sub> | 17.80 (1.00) | 3.98 | 1.14 | NT | 7.83 x 10<sup>8</sup> |
|  | M<sub>16</sub> | EP<sub>14</sub> | 16.80 (2.60) | 6.10 | 2.44 | NT | 4.00 x 10<sup>8</sup> |
|  | M<sub>17</sub> | EP<sub>15</sub> | 18.50 (0.60) | -1.08 | 0.54 | NT | 9.00 x 10<sup>7</sup> |
|  | M<sub>18</sub> | EP<sub>16</sub> | 19.05 (0.70) | 2.08 | 2.08 | NT | 3.83 x 10<sup>8</sup> |
|  | M<sub>19</sub> | EP<sub>17</sub> | 18.40 (0.80) | 2.72 | 0.00 | NT | 4.33 x 10<sup>8</sup> |
|  | M<sub>20</sub> | EP<sub>18</sub> | 19.25 (1.00) | 3.65 | -0.52 | NT | 6.83 x 10<sup>6</sup> |
|  | M<sub>21</sub> | EP<sub>19</sub> | 17.15 (0.80) | 0.00 | -3.39 | NT | 4.17 x 10<sup>7</sup> |
|  | M<sub>22</sub> | EP<sub>20</sub> | 18.60 (0.70) | -0.54 | -0.54 | NT | NA |
| W4 | M<sub>1</sub> | EP<sub>1</sub> | 17.70 (1.40) | 5.08 | 1.69 | OG | 3.33 x 10<sup>9</sup> |
|  | M<sub>2</sub> | EP<sub>2</sub> | 18.00 (1.50) | 7.95 | 5.11 | NT | 5.33 x 10<sup>8</sup> |
|  | M<sub>3</sub> | EP<sub>3</sub> | 20.10 (1.10) | 5.61 | 0.00 | NT | 9.83 x 10<sup>6</sup> |
|  | M<sub>4</sub> | EP<sub>4</sub> | 16.05 (1.00) | 1.84 | -4.29 | NT | 1.30 x 10<sup>9</sup> |
|  | M<sub>5</sub> | EP<sub>5</sub> | 18.10 (0.40) | 1.67 | 1.67 | NT | 8.33 x 10<sup>8</sup> |
|  | M<sub>6</sub> | EP<sub>6</sub> | 18.60 (1.40) | 2.14 | 1.07 | NT | 3.50 x 10<sup>7</sup> |
|  | **M<sub>7</sub>** | **EP<sub>NULL</sub>** | **17.40 (1.00)** | **1.16** | **1.74** | **NI** | **<1.67 x 10<sup>2</sup>** |
|  | M<sub>8</sub> | EP<sub>7</sub> | 17.35 (1.80) | 7.51 | 0.58 | OG | 3.33 x 10<sup>8</sup> |
|  | M<sub>9</sub> | EP<sub>8</sub> | 18.40 (1.80) | 3.19 | -1.06 | NT | 3.83 x 10<sup>8</sup> |
|  | M<sub>10</sub> | EP<sub>9</sub> | 16.70 (1.30) | 5.39 | 1.80 | NT | 1.48 x 10<sup>9</sup> |
|  | M<sub>11</sub> | EP<sub>10</sub> | 17.70 (1.00) | 3.37 | -2.25 | NT | 1.55 x 10<sup>9</sup> |
|  | M<sub>12</sub> | EP<sub>11</sub> | 17.50 (0.90) | 2.27 | -2.84 | NT | 2.83 x 10<sup>8</sup> |
|  | M<sub>13</sub> | EP<sub>12</sub> | 16.70 (0.80) | 2.40 | 0.00 | NT | 1.52 x 10<sup>9</sup> |
|  | M<sub>14</sub> | EP<sub>13</sub> | 20.45 (1.10) | 0.49 | 1.96 | NT | 6.17 x 10<sup>8</sup> |
|  | M<sub>15</sub> | EP<sub>14</sub> | 18.80 (1.30) | 7.14 | 3.85 | NT | 5.33 x 10<sup>8</sup> |
|  | M<sub>16</sub> | EP<sub>15</sub> | 15.65 (1.10) | 6.49 | 2.60 | NT | 2.05 x 10<sup>9</sup> |
|  | M<sub>17</sub> | EP<sub>16</sub> | 17.75 (0.80) | 2.81 | -0.56 | NT | 8.33 x 10<sup>8</sup> |
|  | **M<sub>18</sub>** | **EP<sub>NULL</sub>** | **18.25 (1.40)** | **5.52** | **2.76** | **NI** | **1.00 x 10<sup>3</sup>** |
|  | M<sub>19</sub> | EP<sub>18A</sub> | 19.45 (0.90) | 2.06 | 0.52 | NT* | 1.00 x 10<sup>3</sup> |
|  | M<sub>20</sub> | EP<sub>17</sub> | 16.35 (1.30) | 7.59 | 3.80 | OG | 1.17 x 10<sup>9</sup> |
|  | M<sub>21</sub> | EP<sub>18B</sub> | 18.05 (0.90) | 1.09 | -2.73 | NT | 9.67 x 10<sup>8</sup> |
|  | M<sub>22</sub> | EP<sub>19</sub> | 20.00 (0.20) | -0.50 | -0.50 | NT | NA |
| W5 | M<sub>1</sub> | EP<sub>1</sub> | 18.30 (1.30) | 5.59 | 2.23 | OG | 7.83 x 10<sup>8</sup> |
|  | M<sub>2</sub> | EP<sub>2</sub> | 19.10 (0.90) | 3.70 | 0.53 | YES | 8.50 x 10<sup>7</sup> |
|  | **M<sub>3</sub>** | **EP<sub>NULL</sub>** | **16.40 (1.10)** | **4.27** | **-0.61** | **NI** | **<1.67 x 10<sup>2</sup>** |
|  | M<sub>4</sub> | EP<sub>3</sub> | 18.05 (0.90) | 2.76 | -0.55 | OG | 1.55 x 10<sup>9</sup> |
|  | M<sub>5</sub> | EP<sub>4</sub> | 18.30 (0.70) | 2.20 | 0.55 | NT | 1.58 x 10<sup>9</sup> |
|  | M<sub>6</sub> | EP<sub>5</sub> | 18.50 (1.10) | 4.32 | -1.62 | NT | 1.63 x 10<sup>8</sup> |
|  | M<sub>7</sub> | EP<sub>6</sub> | 16.80 (1.00) | -1.17 | -2.34 | NT | 2.58 x 10<sup>9</sup> |
|  | M<sub>8</sub> | EP<sub>7</sub> | 17.95 (1.70) | 8.47 | 1.69 | NT | 6.17 x 10<sup>8</sup> |
|  | **M<sub>9</sub>** | **EP<sub>NULL</sub>** | **18.40 (1.40)** | **5.43** | **0.54** | **NI** | **3.67 x 10<sup>4</sup>** |
|  | M<sub>10</sub> | EP<sub>8</sub> | 18.60 (1.10) | 2.14 | 2.67 | OG | 3.50 x 10<sup>8</sup> |
|  | M<sub>11</sub> | EP<sub>9</sub> | 18.25 (0.70) | 1.10 | 1.65 | NT | 7.50 x 10<sup>8</sup> |
|  | M<sub>12</sub> | EP<sub>10</sub> | 17.95 (1.70) | 8.94 | 1.12 | NT | 1.98 x 10<sup>9</sup> |
|  | M<sub>13</sub> | EP<sub>11</sub> | 17.15 (1.60) | 8.38 | 3.59 | NT | 1.55 x 10<sup>9</sup> |
|  | M<sub>14</sub> | EP<sub>12</sub> | 18.15 (2.00) | -7.65 | -9.18 | NT | 1.02 x 10<sup>8</sup> |
|  | M<sub>15</sub> | EP<sub>13</sub> | 17.40 (0.80) | 3.45 | 0.00 | NT | 1.00 x 10<sup>9</sup> |
|  | M<sub>16</sub> | EP<sub>14</sub> | 18.00 (0.70) | 2.22 | -1.67 | NT | 9.67 x 10<sup>8</sup> |
|  | M<sub>17</sub> | EP<sub>15</sub> | 17.90 (0.90) | 3.33 | -1.11 | NT | 8.17 x 10<sup>8</sup> |
|  | M<sub>18</sub> | EP<sub>16</sub> | 17.60 (1.10) | 4.57 | -1.71 | NT | 7.33 x 10<sup>8</sup> |
|  | M<sub>19</sub> | EP<sub>17</sub> | 17.50 (0.60) | 3.43 | 2.29 | NT | 2.52 x 10<sup>9</sup> |
|  | M<sub>20</sub> | EP<sub>18</sub> | 18.65 (0.50) | 0.00 | 1.60 | NT | 3.00 x 10<sup>8</sup> |
|  | M<sub>21</sub> | EP<sub>19</sub> | 18.30 (1.20) | 6.70 | 2.79 | NT | 4.67 x 10<sup>8</sup> |
|  | M<sub>22</sub> | EP<sub>20</sub> | 19.50 (1.10) | -5.56 | -5.56 | NT | NA |

## Notes

### Competing Interest Statement

The authors have declared no competing interest.

## References

1. Bull, J. J. Perspective: Virulence. Evolution 48, 1423–1437 (1994).

2. Leventhal, G. E., Hill, A. L., Nowak, M. A. & Bonhoeffer, S. Evolution and emergence of infectious diseases in theoretical and real-world networks. Nat. Commun. 6, 6101 (2015).

3. Hadfield, J. et al. Nextstrain: real-time tracking of pathogen evolution. Bioinformatics 34, 4121–4123 (2018).

4. Ladner, J. T., Grubaugh, N. D., Pybus, O. G. & Andersen, K. G. Precision epidemiology for infectious disease control. Nat. Med. 25, 206–211 (2019).

5. Tomas, M. del M. et al. Hospital outbreak caused by a carbapenem-resistant strain of Acinetobacter baumannii: patient prognosis and risk-factors for colonisation and infection. Clin. Microbiol. Infect. 11, 540–546 (2005).

6. Harris, S. R. et al. Whole-genome sequencing for analysis of an outbreak of methicillin-resistant Staphylococcus aureus: a descriptive study. Lancet Infect. Dis. 13, 130–136 (2013).

7. Worby, C. J., Lipsitch, M. & Hanage, W. P. Shared genomic variants: identification of transmission routes using pathogen deep-sequence data. Am. J. Epidemiol. 186, 1209–1216 (2017).

8. Lieberman, T. D. et al. Genomic diversity in autopsy samples reveals within-host dissemination of HIV-associated M. tuberculosis. Nat. Med. 22, 1470–1474 (2016).

9. Köser, C. U. et al. Rapid whole-genome sequencing for investigation of a neonatal MRSA outbreak. N. Engl. J. Med. 366, 2267–2275 (2012).

10. Andersen, K. G. et al. Clinical sequencing uncovers origins and evolution of Lassa virus. Cell 162, 738–750 (2015).

11. Mate, S. E. et al. Molecular evidence of sexual transmission of Ebola Virus. N. Engl. J. Med. 373, 2448–2454 (2015).

12. Martin, M. A., Lee, R. S., Cowley, L. A., Gardy, J. L. & Hanage, W. P. Within-host Mycobacterium tuberculosis diversity and its utility for inferences of transmission. Microb. Genomics 4, (2018).

13. Lee, R. S., Proulx, J.-F., McIntosh, F., Behr, M. A. & Hanage, W. P. Previously undetected super-spreading of Mycobacterium tuberculosis revealed by deep sequencing. eLife 9, e53245 (2020).

14. Worby, C. J., Lipsitch, M. & Hanage, W. P. Within-host bacterial diversity hinders accurate reconstruction of transmission networks from genomic distance data. PLOS Comput. Biol. 10, e1003549 (2014).

15. Sobel Leonard, A., Weissman, D. B., Greenbaum, B., Ghedin, E. & Koelle, K. Transmission bottleneck size estimation from pathogen deep-sequencing data, with an application to human Influenza A Virus. J. Virol. 91, (2017).

16. Ghafari, M., Lumby, C. K., Weissman, D. B. & Illingworth, C. J. R. Inferring transmission bottleneck size from viral sequence data using a novel haplotype reconstruction method. J. Virol. 94, (2020).

17. Read, H. M. et al. The in vitro and in vivo effects of constitutive light expression on a bioluminescent strain of the mouse enteropathogen Citrobacter rodentium. PeerJ 4, e2130 (2016).

18. Wiles, S. et al. Organ specificity, colonization and clearance dynamics in vivo following oral challenges with the murine pathogen Citrobacter rodentium. Cell. Microbiol. 6, 963–972 (2004).

19. Wiles, S., Pickard, K. M., Peng, K., MacDonald, T. T. & Frankel, G. In vivo bioluminescence imaging of the murine pathogen Citrobacter rodentium. Infect. Immun. 74, 5391–5396 (2006).

20. Mullineaux-Sanders, C. et al. Citrobacter rodentium relies on commensals for colonization of the colonic mucosa. Cell Rep. 21, 3381–3389 (2017).

21. Bishop, A. L., Wiles, S., Dougan, G. & Frankel, G. Cell attachment properties and infectivity of host-adapted and environmentally adapted Citrobacter rodentium. Microbes Infect. 9, 1316–1324 (2007).

22. Wiles, S., Dougan, G. & Frankel, G. Emergence of a ‘hyperinfectious’ bacterial state after passage of Citrobacter rodentium through the host gastrointestinal tract. Cell. Microbiol. 7, 1163–1172 (2005).

23. Lythgoe, K. A. et al. SARS-CoV-2 within-host diversity and transmission. Science (2021) doi:10.1126/science.abg0821.

24. Lieberman, T. D. et al. Genetic variation of a bacterial pathogen within individuals with cystic fibrosis provides a record of selective pressures. Nat. Genet. 46, 82–87 (2014).

25. Hall, M. D. et al. Improved characterisation of MRSA transmission using within-host bacterial sequence diversity. eLife 8, e46402 (2019).

26. Didelot, X., Fraser, C., Gardy, J. & Colijn, C. Genomic infectious disease epidemiology in partially sampled and ongoing outbreaks. Mol. Biol. Evol. 34, 997–1007 (2017).

27. Wymant, C. et al. PHYLOSCANNER: Inferring Transmission from Within- and Between-Host Pathogen Genetic Diversity. Mol. Biol. Evol. 35, 719–733 (2018).

28. Brown, C. M. Outbreak of SARS-CoV-2 infections, including COVID-19 vaccine breakthrough infections, associated with large public gatherings — Barnstable County, Massachusetts, July 2021. MMWR Morb. Mortal. Wkly. Rep. 70, (2021).

29. Lemieux, J. E. et al. Phylogenetic analysis of SARS-CoV-2 in Boston highlights the impact of superspreading events. Science 371, eabe3261 (2021).

30. Read, H., Patel, P. & Wiles, S. Initiating and monitoring natural infection of mice by bioluminescent Citrobacter rodentium. protocols.io https://www.protocols.io/view/initiating-and-monitoring-natural-infection-of-mic-b9r9r596 (2022).

31. Bates, D., Mächler, M., Bolker, B. & Walker, S. Fitting Linear Mixed-Effects Models using lme4. J. Stat. Softw. 67, 1–48 (2015).

32. Li, H. & Durbin, R. Fast and accurate short read alignment with Burrows–Wheeler transform. Bioinformatics 25, 1754–1760 (2009).

33. Li, H. et al. The Sequence Alignment/Map format and SAMtools. Bioinforma. Oxf. Engl. 25, 2078–2079 (2009).

34. Poplin, R. et al. Scaling accurate genetic variant discovery to tens of thousands of samples. http://biorxiv.org/lookup/doi/10.1101/201178 (2017) doi:10.1101/201178.

35. Quinlan, A. R. & Hall, I. M. BEDTools: a flexible suite of utilities for comparing genomic features. Bioinformatics 26, 841–842 (2010).

36. Magaziner, S. J., Zeng, Z., Chen, B. & Salmond, G. P. C. The prophages of Citrobacter rodentium represent a conserved family of horizontally acquired mobile genetic elements associated with enteric evolution towards pathogenicity. J. Bacteriol. 201, (2019).

37. Danecek, P. et al. The variant call format and VCFtools. Bioinformatics 27, 2156–2158 (2011).

38. Katoh, K. & Standley, D. M. MAFFT Multiple Sequence Alignment Software Version 7: Improvements in performance and usability. Mol. Biol. Evol. 30, 772–780 (2013).

39. Stamatakis, A. RAxML version 8: a tool for phylogenetic analysis and post-analysis of large phylogenies. Bioinformatics 30, 1312–1313 (2014).

40. Kumar, S., Stecher, G. & Tamura, K. MEGA7: Molecular Evolutionary Genetics Analysis Version 7.0 for Bigger Datasets. Mol. Biol. Evol. 33, 1870–1874 (2016).

